# Cortical tension as a mechanical barrier to safeguard against premature differentiation during neurogenesis

**DOI:** 10.1101/2025.09.19.677444

**Authors:** Daniel Halperin, Chrystian Junqueira Alves, Marcelle Rodrigues Lemos, Swagata Dey, Haochen Tao, Sangjo Kang, Xianting Li, Zhenyu Yue, Jiaxi Li, Rodrigo Alves Dias, Gustavo de Oliveira Rosa, José P. Rodrigues Furtado de Mendonça, Molly Estill, Yonatan Perez, Susana I. Ramos, Nadejda M. Tsankova, Li Shen, Roland H. Friedel, Hongyan Zou

## Abstract

Neuronal differentiation requires coordinated gene reprogramming and morphodynamic remodeling. How mechanical forces integrate with nuclear gene programs during neurogenesis remains unresolved. Here, we identify cortical tension as a mechanical barrier that safeguards against premature neuronal differentiation. Deletion of Plexin-B2, a guidance receptor controlling actomyosin contractility, lowers this barrier, enabling neurite outgrowth and accelerating neuronal lineage commitment. We show that coupling of extrinsic differentiation cues with intrinsic morphodynamics is essential for stabilizing neuronal fate and that cortical barrier and epigenetic barrier act in concert to regulate developmental timing. In cerebral organoids, Plexin-B2 ablation triggered premature cell-cycle exit and differentiation, resulting in progenitor pool depletion and neuroepithelial disorganization, phenotypes echoing intellectual disability in patients with rare pathogenic *PLXNB2* variants. Our studies demonstrate that cortical tension functions as mechano-checkpoint that regulates the onset of neurogenesis. Lowering this barrier may provide a strategy to accelerate induced neuron generation and maturation for CNS disease modeling.

## INTRODUCTION

Neurogenesis has long been viewed through the lens of gene regulatory programs. Neuronal differentiation requires precise spatial and temporal control of cell-cycle exit and activation of lineage-specific transcriptional programs ^1,2^. Extensive work has defined the growth factors, transcriptional regulators, and gene networks that govern the transition from neural progenitor cells (NPCs) to differentiating neurons ^3–7^. More recently, an “epigenetic barrier” has been proposed to safeguard the timing of transcriptional maturation and prevent premature differentiation ^8^.

Neuronal lineage commitment also requires major biomechanical remodeling to enable neurite protrusion. While cytoskeletal dynamics have been implicated in early neuronal morphogenesis ^9^, how biomechanical forces are integrated with transcriptional and epigenetic programs remains poorly understood. A related question concerns the role of cell mechanics of NPCs in neuroepithelial morphogenesis, particularly during neural tube closure, a crucial process during neurodevelopment ^10^.

Plexin-B2, an evolutionarily conserved axon guidance receptor, has emerged as a regulator of actomyosin contractility and cell stiffness in human embryonic stem cells (hESCs) and NPCs ^11^. In vivo, *Plxnb2* knockout mice display neural tube closure defects ^12^, and combined deletion of *Plxnb1/2* in the CNS leads to cortical thinning as a result of dysregulated NPC differentiation and cell-cycle exit ^13^. Clinically, recent human genetic studies have identified rare pathogenic *PLXNB2* variants in families with intellectual disability ^14^, highlighting its relevance to human brain development.

Here, we use induced neuron and cerebral organoid models to uncover a previously unrecognized role of cortical tension as a mechanical barrier that safeguards against premature neuronal differentiation. We show that Plexin-B2 fortifies the cortical actin network to delay neurite initiation. Loss of Plexin-B2 lowers this barrier, resulting in early neurite protrusion, accelerated neuronal differentiation, and transcriptional signatures of neuronal lineage commitment. We further demonstrate that coupling of extrinsic differentiation cues with intrinsic biomechanical states is required to stabilize neuronal fate, and that cortical mechanical barrier operates in concert with epigenetic barrier to maintain developmental timing.

Finally, cerebral organoid studies reveal that Plexin-B2 deficiency causes premature NPC cell-cycle exit, progenitor pool depletion, and disruption of neuroepithelial integrity. Structure-function analysis shows that Plexin-B2 signals via its Ras-GAP domain and depends on its flexible extracellular ring domain to regulate cortical mechanics. Together, these findings identify cortical tension as a mechano-checkpoint for neurogenesis, thereby providing new insight into neurodevelopmental disorders and offering strategies to accelerate induced neuron generation for disease modeling.

## RESULTS

### Decline of cortical actin tension during neuronal differentiation

To investigate how neuronal differentiation is coupled to morphodynamic remodeling, we examined cortical filamentous (F-) actin during the derivation of induced neurons (iNs) from human embryonic stem cells (hESCs) using a dual-SMAD inhibition protocol ^15^ (**Fig. 1a**). Phalloidin staining revealed a progressive reduction in cortical F-actin during neuroinduction and differentiation, coinciding with an upregulation of the neuronal marker β3-tubulin (TUJ1) (**Fig. 1b, d**). Consistent with reduced cortical tension, phosphorylated myosin light chain 2 (pMLC2), a marker of actomyosin contractility, declined markedly from the NPC stage at day (D) 10 to fully iN stage at D36 (**Fig. 1c**). Together, these data show that neuronal differentiation is accompanied by a major cytoskeletal reorganization, with progressive lowering of cortical tension linked to neurite formation.

**Figure 1.**
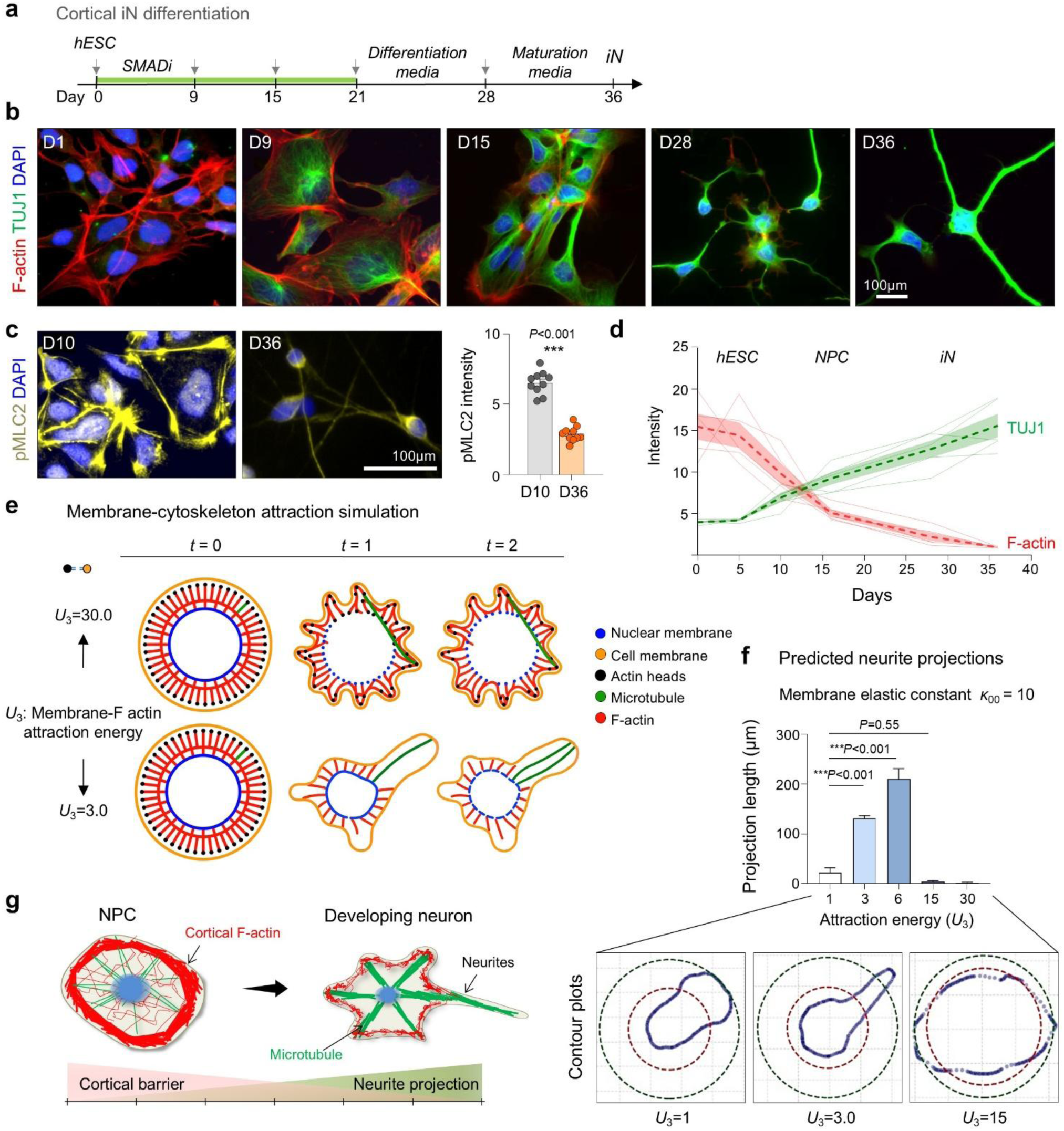
Progressive decline of cortical actin tension during neuronal differentiation. **a)** Schematic of the dual-SMAD inhibition (SMADi) protocol to generate induced neurons. **b-d)** Progressive decrease of F-actin and pMLC2 and reciprocal increase of TUJ1 expression along iN differentiation process. Graphs represent mean ± SEM; n = 3 cultures for each condition; two-tailed nested t-test. e) Molecular dynamics simulation of neurite morphogenesis with varying membrane-actin attraction energy (*U*_3_; high = 30.0 vs. low = 3.0) and fixed membrane tension (*κ*_00_ = 500). Lower *U*_3_ facilitated dynamic protrusions resembling neurite extension. f) Quantification of simulated neurite projection length as a function of *U_3_*, with *κ*_00_ fixed at 10. Bar graphs represent mean ± SEM, one-way ANOVA with Dunnett’s multiple-comparison test. Intermediate *U_3_* values promoted maximal extension, while very high or low *U*_3_ restricted membrane deformation and neurite protrusions. Contour plots below show representative simulated cell morphologies. g) Model: Cortical actin functions as a structural barrier restricting premature neurite projections.

To dissect the underlying biophysical principles, we performed molecular dynamics simulations using a coarse-grained bead-spring model representing the plasma membrane, F-actin, microtubules, and nuclear envelope (**Fig. 1e**). Distinct morphodynamic outcomes emerged depending on the balance of membrane-actin attraction energy (*U*_3_; modeling cortical tension) and membrane elastic constant (*κ*_00_; modeling membrane tension). Moderate *U*_3_ values supported stable neurite-like protrusions, whereas excessively high or low *U*_3_ values impaired projection stability (**Fig. 1e, f**; **Fig. S1a-c**). Mathematical simulations thus highlight the requirement for balanced cortical and membrane tension forces to permit robust neurite initiation during neuronal development (**Fig. 1g**).

### Lowering cortical tension enables early neurite protrusions

We next tested whether cortical tension acts as a barrier to premature neurite initiation. Our prior work showed that Plexin-B2 regulates actomyosin contractility in hESCs and NPCs via Rap GTPases ^11^. We therefore tested whether Plexin-B2-mediated cortical tension influences the developmental timing of neurite protrusions by comparing neuroinduction of WT and *PLXNB2^-/-^*hESCs (**Fig. 2a**). Plexin-B2 deletion did not affect pluripotency marker expression (NANOG, OCT4, SOX2) (**Fig. S2a**).

**Figure 2.**
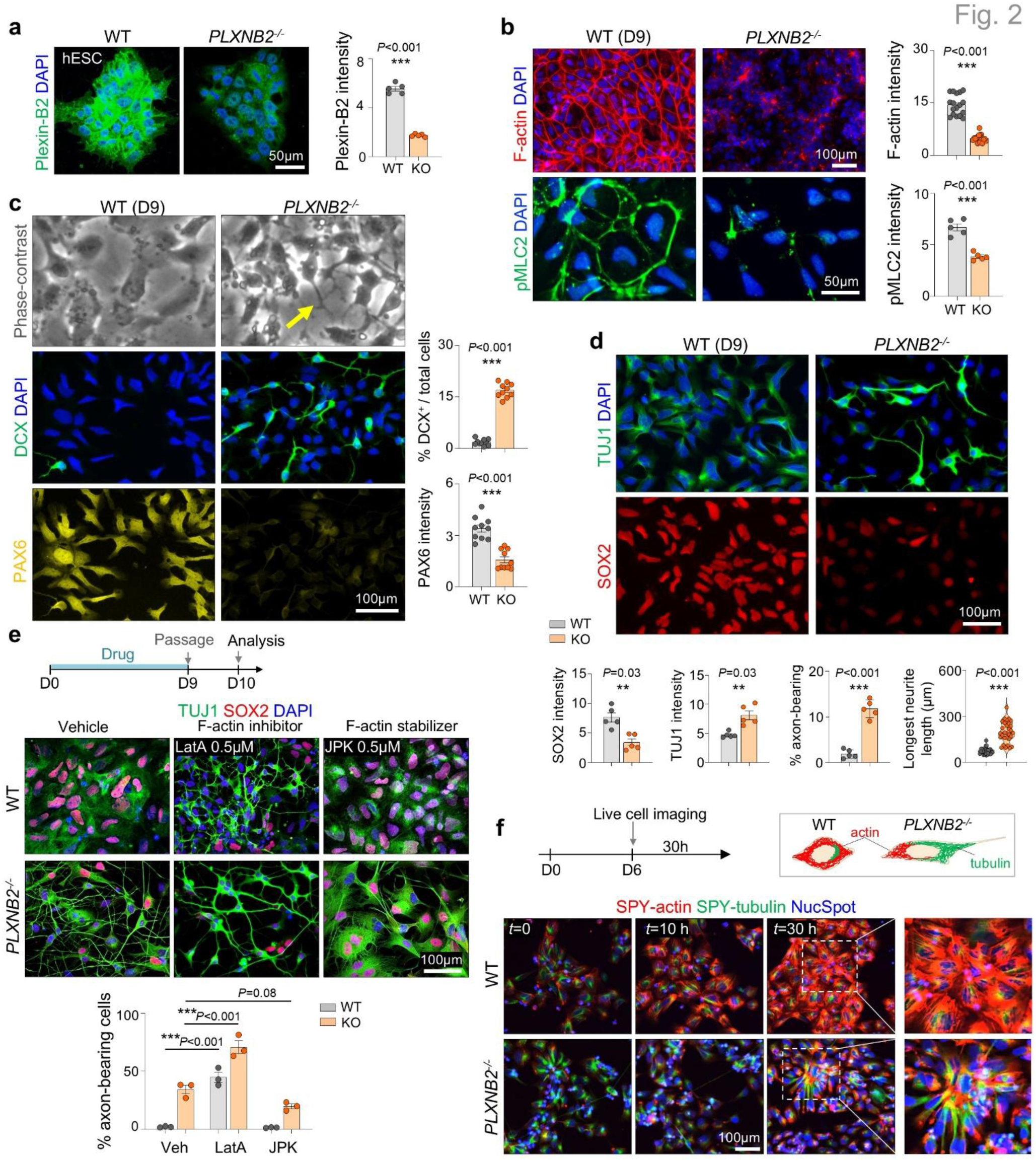
Lowering cortical tension accelerates neurite formation. **a)** IF showing loss of Plexin-B2 protein in *PLXNB2^⁻/⁻^* hESCs. n = 5 independent cultures per condition; unpaired two-tailed t-test. Bar graphs represent mean ± SEM. **b)** At D9 of differentiation, *PLXNB2^⁻/⁻^* cells displayed reduced cortical F-actin and pMLC2 compared with WT. n = 5 independent cultures per condition, two-tailed nested t-test. **c)** Phase-contrast (top) and IF (bottom) images at D9. *PLXNB2^⁻/⁻^*cells formed elongated projections (arrows) and showed increased DCX with reduced PAX6 compared with WT. n *=* 10 fields across two independent cultures; unpaired two-tailed t-test. **d)** IF for TUJ1 and SOX2 at D9. *PLXNB2^⁻/⁻^* cells showed increased TUJ1 and reduced SOX2. Each data point represents the mean of multiple fields of view from two independent cultures; two-tailed nested t-test. Bar graphs represent mean ± SEM. **e)** Schematic and representative IF images from epistasis analysis. Latrunculin A (LatA, 0.5 µM) reduced cortical F-actin and promoted neurite protrusions in WT cells, mimicking Plexin-B2 knockout. Conversely, jasplakinolide (JPK, 0.5 µM) stabilized F-actin and suppressed projections in *PLXNB2^⁻/⁻^* cells, restoring SOX2 expression. n = 3 images per condition; one-way ANOVA with Tukey’s test. Bar graphs represent mean ± SEM. **f)** Live-cell imaging of D6 cells labeled with NucSpot, SPY-tubulin, and SPY-actin over 30 hours. WT cells progressively reinforced cortical F-actin without protrusions, whereas *PLXNB2^-/-^* cells showed diminished cortical actin and long tubulin-based projections.

At day 9, *PLXNB2^-/-^* cells displayed reduced cortical F-actin and pMLC2 compared with WT (**Fig. 2b**), consistent with prior observations in hESCs ^11^. Strikingly, *PLXNB2^-/-^*cells extended elongated neurite-like protrusions, whereas WT cells retained a rounded morphology without long processes (**Fig. 2c**). Immunofluorescence (IF) revealed premature induction of neuronal markers (DCX, TUJ1) and MeCP2, a neuronal maturation marker ^16^, with concomitant reduction of progenitor markers (SOX2, PAX6, Nestin) and proliferative marker Ki67, consistent with accelerated neuronal differentiation (**Fig. 2c, d**; **Fig. S2b, c**).

Epistasis analyses confirmed a mechanical basis for this phenotype. In WT cells, actin depolymerization with low dose of latrunculin A promoted neurite outgrowth and downregulated SOX2, phenocopying *PLXNB2* knockout. Conversely, stabilizing F-actin with jasplakinolide suppressed neurite projections in *PLXNB2^-/-^* cells and restored SOX2 expression (**Fig. 2e**).

Live-cell imaging of D6 NPCs labeled with SPY-Actin, SPY-Tubulin, and NucSpot during induction revealed progressive cortical actin assembly without protrusions in WT NPCs, whereas *PLXNB2^-/-^* NPCs showed reduced cortical actin but long tubulin-containing neurites (**Fig. 2f**; **Movie S1**). Together, these findings identify cortical actin as a mechanical barrier to premature neurite initiation. Lowering this barrier either pharmacologically or via Plexin-B2 deletion promotes early neurite initiation and accelerates neuronal lineage commitment.

### Gene expression analysis confirms accelerated neuronal cell fate acquisition of *PLXNB2^-/-^* cells

To validate accelerated neuronal differentiation in *PLXNB2^-/-^*(KO) cells, we performed single-nucleus RNA-seq (snRNA-seq) at D9 of neuroinduction (n = 3 per genotype), yielding 7,800 high-quality nuclei (**Fig. 3a**). UMAP analysis showed clear segregation of WT (n = 4,974) and KO (n = 2,826) cells, with WT enriched for progenitor markers (*SOX2, NES*), and KO enriched for neuronal markers (*DCX, TUBB3*) (**Fig. 3a, b**).

**Figure 3.**
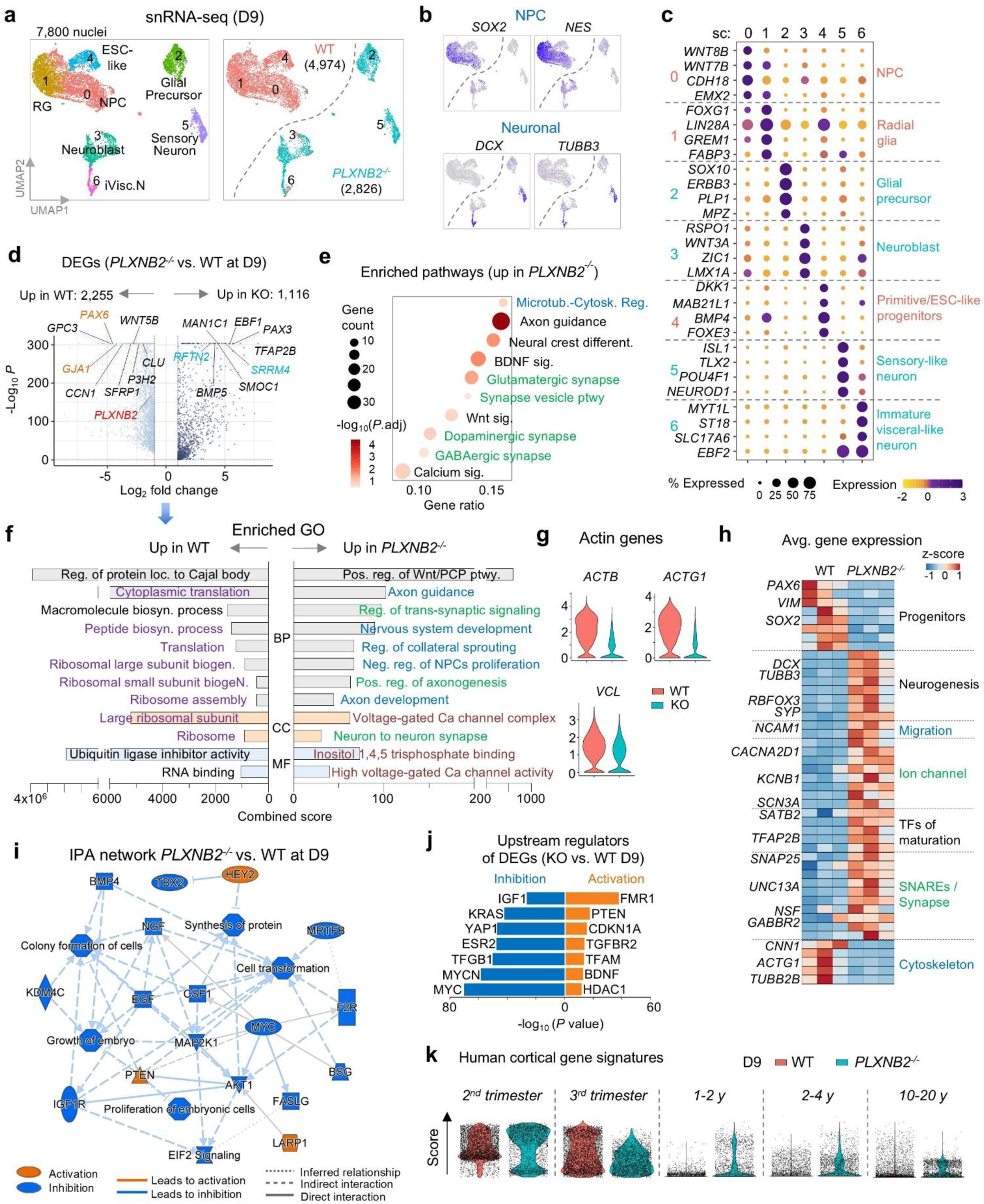
Single-nucleus transcriptomics confirms accelerated neuronal differentiation of *PLXNB2⁻^/^⁻* cells. **a)** UMAP embedding of snRNA-seq data from D9 cells (n = 3 replicates per genotype). WT cells segregated into clusters enriched for radial glia (RG), neural progenitors (NPCs), and ESC-like states, whereas KO cells aligned with differentiated neuronal clusters. **b)** Feature plots showing expression of NPC and neuronal markers. **c)** Expression of representative marker genes across D9 subclusters. **d)** Volcano plot of differentially expressed genes (DEGs; *PLXNB2*^-/-^ vs. WT), with selected genes highlighted. **e)** Bubble plot of enriched pathways upregulated in D9 *PLXNB2*^⁻/⁻^ cells. **f)** ENRICHR GO enrichment analysis of DEGs for categories of biological process (BP), cellular component (CC), and molecular function (MF), color-coded by theme. **g)** Violin plots showing downregulation of actin cytoskeleton–associated genes in *PLXNB2^-/-^* cells. **h)** Heatmap of DEGs grouped by functional categories, across three replicates per genotype. **i)** Ingenuity Pathway Analysis (IPA) network summary. Growth factor signaling pathways were broadly suppressed, while PTEN was activated in *PLXNB2*^⁻/⁻^ cells. **j)** Predicted upstream regulators (IPA) of DEGs in *PLXNB2*^⁻/⁻^vs. WT cells. **k)** Developmental stage scoring against human cortex gene signatures (Velmeshev et al., 2023). D9 WT cells aligned with 2^nd^-3^rd^ trimester profiles, whereas *PLXNB2*^⁻/⁻^ cells shifted toward postnatal signatures.

Subclustering revealed that WT cells consisted largely of progenitors and radial glia, whereas KO cells partitioned into neuronal and neuroblast subtypes (**Fig. 3c**). Differential expression analysis identified over 3,300 genes significantly altered in KO cells (adj. *P*<0.05, log_2_FC>1), with upregulation of neuronal, ion channel, and synaptic programs, and downregulation of progenitor and cytoskeletal regulators (**Fig. 3d**). Gene ontology (GO) enrichment confirmed activation of pathways related to neurogenesis, axonogenesis, and synaptic signaling, while biosynthetic and ribosomal processes were suppressed in KO cells (**Fig. 3e-g**). Heatmap analysis further highlighted the shift from progenitor to neuronal transcriptional programs (**Fig. 3h**).

Ingenuity network summary analysis highlighted global inhibition of growth factor pathways (NGF, BMP, EGF, IGF, MAP2K1, CSF1) in KO cells, with activation of only PTEN, HEY2 (a notch target gene important for neurodevelopment ^17^, and LARP1 (linked to Autism ^18^) (**Fig. 3i**). Upstream regulator analysis predicted activation of neuronal differentiation drivers and cell-cycle inhibitors, alongside inhibition of proliferative and growth factor pathways (**Fig. 3j**). This included activation of FMR1 (Fragile X Mental Retardation 1), an RNA binding protein required for cortical development ^19^), PTEN (regulator of NPC proliferation and self-renewal ^20,21^), CDKN1A/p21 (cell cycle inhibitor), and BDNF (neurotrophic factor). In contrast, MYC, MYCN, KRAS, YAP1 and IGF (a growth factor for embryonic stem cell renewal ^22^) were inhibited.

Finally, developmental signature mapping using human brain reference atlas ^7^ revealed that D9 WT cells mapped predominantly to prenatal (2^nd^ or 3^rd^ trimester) profiles, whereas KO cells also aligned with postnatal signatures (**Fig. 3k**). Together, these data show that Plexin-B2 loss lowers cortical actin barriers, leading to precocious neuronal lineage commitment and maturation.

### Function-blocking nanobodies against Plexin-B2 accelerate neuronal differentiation

To further test whether Plexin-B2 activity restrains neuronal differentiation, we generated function-blocking nanobodies (Nbs) against human Plexin-B2 extracellular domain. Camelid immunization followed by phage display yielded twenty high-affinity VHH domains, two of which were expressed as VHH-Fc fusions for further testing (**Fig. 4a**). Continuous Nb treatment during neuroinduction weakened the cortical F-actin ring and induced early neurite protrusions, while control Nb (raised against an unrelated antigen) had no effect (**Fig. 4b-d**). Anti-Plexin-B2 Nbs also increased expression of DCX and TUJ1 while reducing SOX2, recapitulating the phenotype of *PLXNB2^-/-^*cells. These findings provide independent confirmation that Plexin-B2 functions as a checkpoint, preventing premature neuronal differentiation.

**Figure 4.**
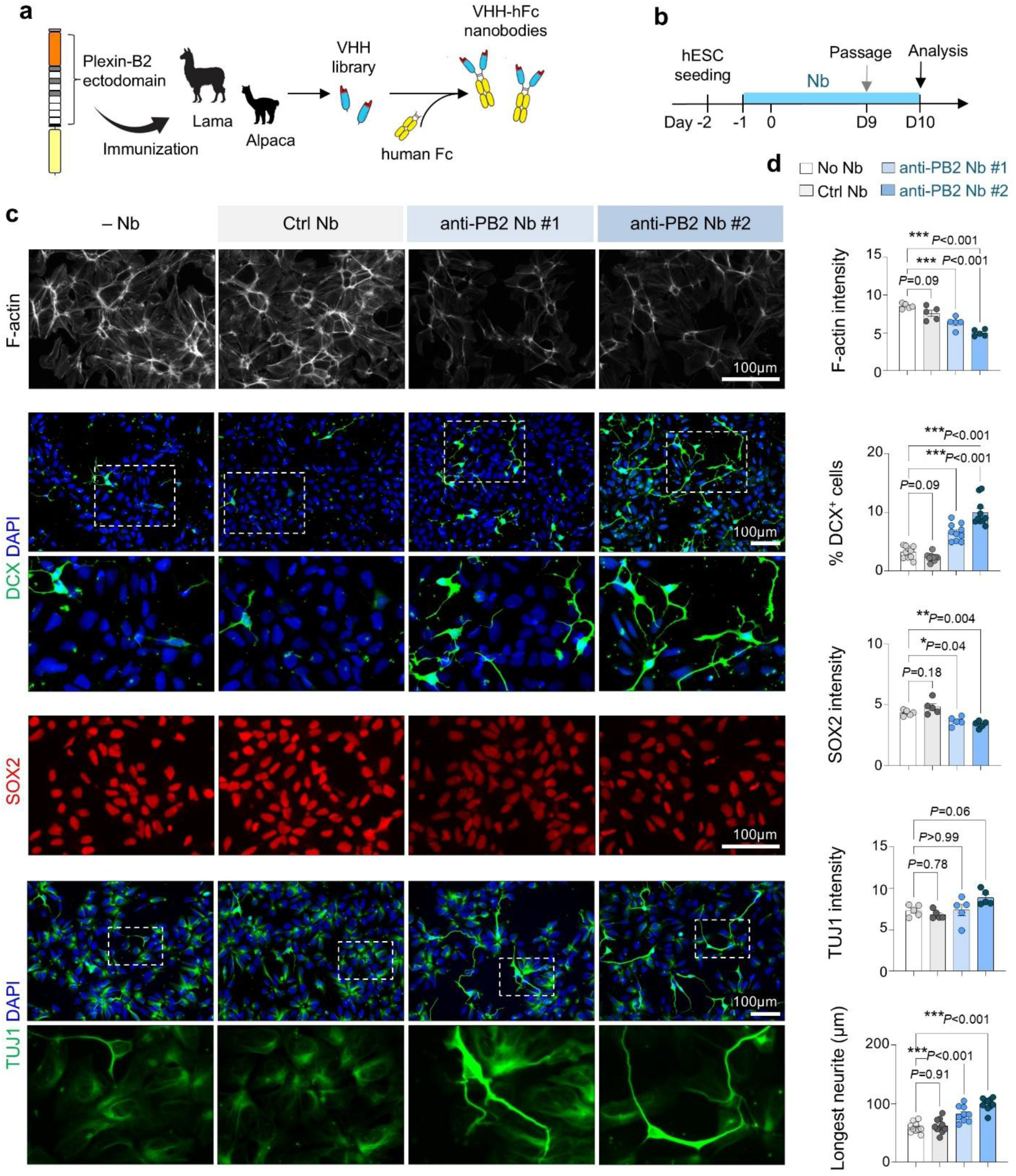
Function-blocking nanobodies against Plexin-B2 accelerate neuronal differentiation. **a)** Workflow for generating anti-Plexin-B2 (PB2) nanobodies (Nbs). Camelids were immunized with recombinant extracellular domain of human PB2 protein, followed by phage display to isolate high-affinity VHHs, which were fused to human Fc (hFc). **b)** Experimental design. VHH-Fc Nbs (10 μg/ml) were added one day before iN protocol. **c)** Representative images showing reduced cortical F-actin, increased DCX and TUJ1, and decreased SOX2 in D10 cells treated with two independent anti-PB2 Nbs compared with control or no Nb. **d)** Quantification of marker expression. Each dot represents the mean of a field of view. n = 5 cultures for each condition; one-way ANOVA with Dunnett’s multiple-comparison test. Bar graphs represent mean ± SEM.

### Timed maturation cues stabilize neuronal identity of *PLXNB2*^-/-^ induced neurons

We next asked whether the early neuronal identity of *PLXNB2^-/-^*cells could be stably maintained. We found that at later stages (D28 and D36) under either forebrain (FB) or midbrain (MB) protocols, whereas WT cells developed into expected iN phenotypes, *PLXNB2^-/-^* cells lost TUJ1⁺ neurites and adopted a flattened morphology, suggesting instability of neuronal identity (**Fig. 5a; Fig. S3a-e**).

**Figure 5.**
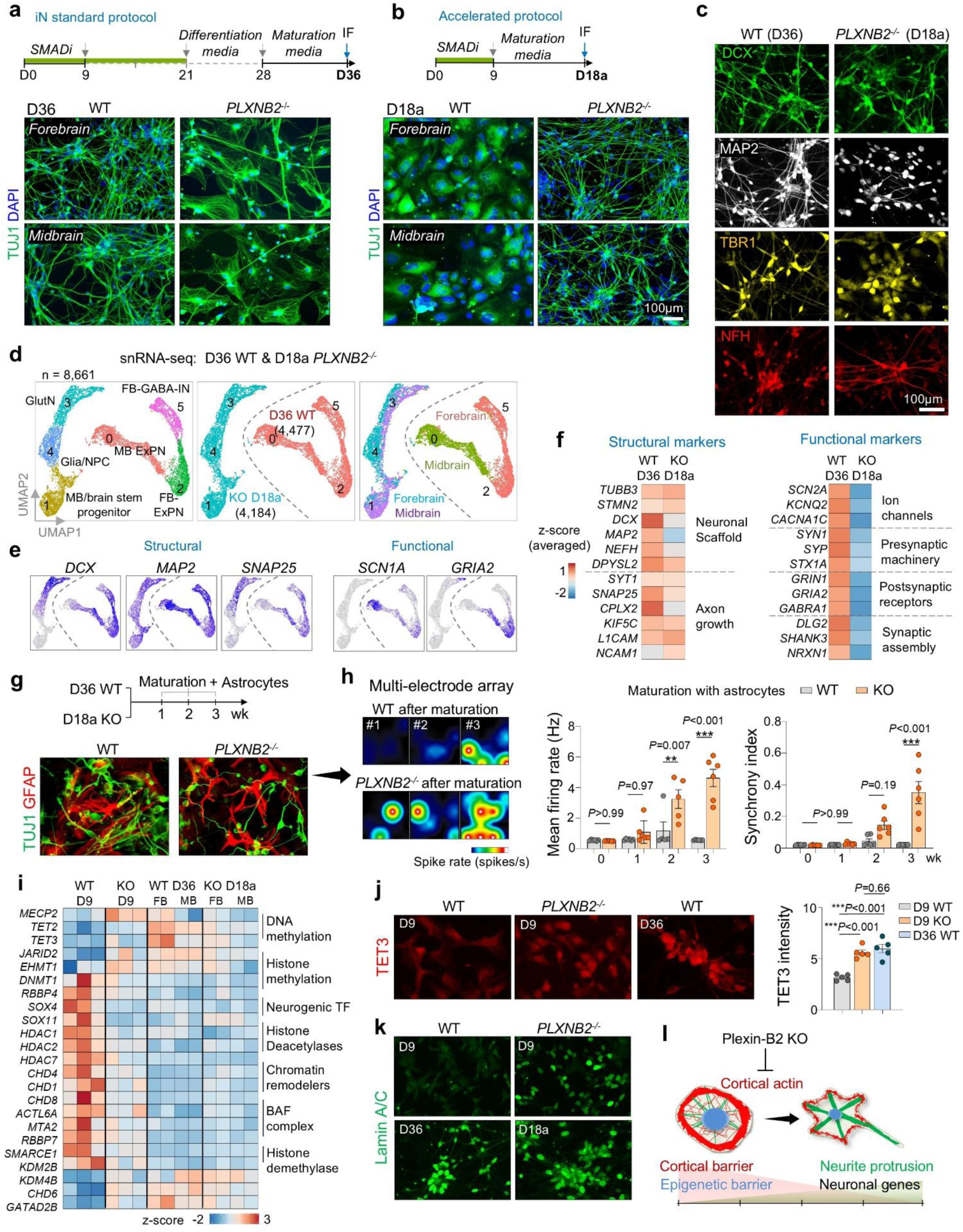
Matching external cues with intrinsic morphodynamics safeguards neuronal lineage commitment. **a)** Standard forebrain and midbrain neuronal differentiation protocols. At D36, WT cells generated dense networks of TUJ1⁺ axon-bearing neurons, whereas *PLXNB2^⁻/⁻^* cells displayed aberrant morphologies. **b)** Accelerated protocol. After passage at D9, cells were switched directly to maturation medium, bypassing the differentiation media step. By D18a, *PLXNB2^⁻/⁻^* cells developed dense networks of TUJ1⁺ axon-bearing neurons, while WT cells retained progenitor-like morphology. **c)** IF comparison of neuronal markers and morphology in D36 WT (standard protocol) and D18a *PLXNB2^⁻/⁻^* (accelerated protocol) cells. **d)** UMAP embedding of D18a *PLXNB2⁻/⁻* and D36 WT iNs, showing segregation by genotype and forebrain vs. midbrain protocols. **e)** Feature plots showing shared neuronal markers across subclusters and genotypes. **f)** Transcriptional profiling revealed comparable expression of core neuronal/axonal markers between D36 WT and D18a *PLXNB2^⁻/⁻^* cells, but lower expression of functional genes in the latter. **g)** Experimental timeline and IF images show co-culture of iNs with GFAP⁺ astrocyte lawns to promote maturation. **h)** Multi-electrode array (MEA) recordings. *PLXNB2^⁻/⁻^* cells after maturation exhibited higher firing rates and synchrony index than WT cells (n = 6 cultures per condition; one-way ANOVA with Dunnett’s correction). Bar graphs represent mean ± SEM. **i)** Heatmap of epigenetic regulators shows convergent transcriptional shifts of D36 WT and D18a *PLXNB2^⁻/⁻^* iNs relative to D9 WT cells. D9 *PLXNB2^⁻/⁻^* cells already exhibited a similar shift, indicating accelerated epigenetic reprogramming. **j)** D9 *PLXNB2^⁻/⁻^* cells showed elevated nuclear TET3 compared with D9 WT, reaching levels comparable to D36 WT. n = 5 fields from 2 independent experiments for each condition, nested one-way ANOVA with Tukey’s multiple-comparison test. **k)** D9 *PLXNB2^⁻/⁻^* cells exhibited precocious expression of lamin A/C, similar to D36 WT and D18a *PLXNB2^⁻/⁻^* iNs. **l)** Working model: Plexin-B2-mediated cortical tension act as a mechanical barrier alongside an epigenetic barrier to prevent precocious differentiation.

snRNA-seq revealed that D36 WT cells expressed neuronal markers (*DCX, RORB, RBFOX3, TUBB3*), whereas KO cells shifted toward mesenchymal, fibrotic and stress-associated states enriched for ECM genes (*FN1, COL1A1*) (**Fig. S4a, b**). Subcluster analysis confirmed WT cells generated forebrain and midbrain excitatory neurons and interneurons, while KO cells comprised mesenchymal/EMT-like populations (**Fig. S4c, d**). Differential expression showed downregulation of axon guidance and synaptic genes in KO cells and upregulation of ECM and senescence pathways (**Fig. S4e-h**). Thus, mismatched cues destabilize neuronal lineage commitment in *PLXNB2^-/-^*cells.

To test whether premature neuronal differentiation could be stabilized, we advanced the maturation media step to D9 (instead of D28). With this accelerated protocol, *PLXNB2^-/-^* cells maintained robust TUJ1⁺ networks and expressed neuronal markers (DCX, MAP2, TBR1, NFH), while WT cells remained progenitor-like when exposed prematurely to maturation cues (**Fig. 5b, c**). Transcriptomic profiling of cells at day 18 of the accelerated protocol (D18a) showed that *PLXNB2^-/-^* cells adopted defined forebrain and midbrain neuronal subclusters with strong expression of neuronal and synaptic genes (*DCX, RBFOX3, SNAP25*), whereas D18a WT cells retained progenitor signatures (*PAX6, MKI67*) and undefined progenitor state (**Fig. S5a-d)**.Pathway and upstream regulator analyses confirmed activation of neuronal differentiation programs (BDNF, NGF, ASCL1) and suppression of progenitor regulators (REST, EZH2, YAP1) in KO cells (**Fig. S5e-j**). By contrast, D18a WT cells due to premature exposure to maturation cues upregulated genes for ribosome biogenesis, ubiquitin ligase activity, proliferation, and focal adhesion, signifying stalled neuronal progression.

Together, these data demonstrate that temporally matched maturation cues are required to stabilize the accelerated neuronal differentiation of *PLXNB2^-/-^* cells and prevent lineage instability.

### Functional status of *PLXNB2^-/-^* neurons after accelerated differentiation

To assess functionality, we further compared D18a *PLXNB2^-/-^*iNs with D36 WT iNs. snRNA-seq integration showed adjacent clustering according to genotype and differentiation protocol (**Fig. 5d**). WT cells formed forebrain excitatory neurons, GABAergic interneurons, and midbrain excitatory neurons, while KO cells comprised maturing glutamatergic neurons, glia-neuronal precursors, and brainstem progenitors (**Fig. 5d**). Both groups expressed neuronal and axonal markers, but *PLXNB2^-/-^* iNs expressed lower levels of voltage-gated ion channels (*KCNB1, KCNC1, CACNA1D*) and synaptic receptors (*GABRB3, GRIN1/2*), a sign of delayed acquisition of synaptic programs at D18a compared to WT iNs at D36 (**Fig. 5e, f**). Notably, while some of these genes are broadly involved in excitatory neurotransmission (*KCNB1, CACNA1D, GRIN1/2*), other genes (*KCNC1* and *GABRB3)* are linked to inhibitory function.

We further matured iNs by co-culturing with astrocytes for 1-3 weeks followed by neuronal activity evaluation. After co-culturing with astrocytes, both WT and KO iNs formed dense TUJ1⁺ neuronal networks alongside GFAP⁺ astrocytes (**Fig. 5g**). Multi-electrode recordings revealed that by 2-3 weeks of maturation with astrocytes, KO iNs displayed enhanced firing rates and greater network synchrony compared to WT cells, indicating that they can undergo rapid functional maturation under appropriate environmental cues.

### Cytoskeletal barrier is coupled with epigenetic barrier

We next asked whether the cortical mechanical barrier intersects with the epigenetic barrier that safeguards progenitor identity ^8^. Gene expression analysis showed that chromatin regulator profiles of D18a KO iNs resembled those of D36 WT iNs but were distinct from D9 WT progenitors (**Fig. 5i**). Notably, D9 KO cells already expressed epigenetic signatures characteristic of D36 WT iNs, suggesting accelerated epigenetic reprogramming with lowered cortical barrier.

For further verification, we examined the expression of Tet3, an epigenetic regulator of DNA hydroxymethylation that is upregulated in neuronal differentiation ^23^. IF confirmed elevated Tet3 in D9 KO cells, comparable to D36 WT neurons (**Fig. 5j**). Nuclear remodeling was also accelerated: the levels of Lamin A/C, an essential nuclear envelope component involved in chromatin remodeling and neuronal lineage commitment ^24^, were higher in D9 KO cells and reached mature levels in D18a KO iNs, paralleling D36 WT iNs (**Fig. 5k**). Since cytoskeletal forces are transmitted to the nucleus to influence chromatin states ^25,26^, these findings support a mechanistic coupling of cortical mechanics and epigenetic programming.

Together, these results identify mechano-epigenetic checkpoints that synchronize cytoskeletal tension, nuclear remodeling, and transcriptional reprogramming to ensure fidelity of neuronal lineage commitment and neurite morphogenesis (**Fig. 5l).**

### Plexin-B2 safeguards progenitor pool integrity and corticogenesis in cerebral organoids

Human genetic studies have linked biallelic *PLXNB2* variants to intellectual disability ^14^. To test whether Plexin-B2 functions as a mechanical checkpoint during human corticogenesis, we generated 3D cerebral organoids from WT and *PLXNB2^-/-^* hESCs using the Lancaster protocol ^27^ (**Fig. S6a**). By day 42, WT organoids developed ventricle-like structures lined by SOX2⁺ progenitors and cortical plates containing newborn neurons, with Plexin-B2 broadly expressed in ventricular zones, cortical plate, and outer radial glia, a primate-specific progenitor population critical for cortical expansion (**Fig. 6a, b**). Similar patterns were observed in human fetal cortex at 23 weeks (**Fig. 6b**). The Plexin-B2 ligands SEMA4B and SEMA4C were also detected in cortical structures (**Fig. S6b**). Analysis of human cortical single-cell transcriptomics ^7^ confirmed broad *PLXNB2* expression across developmental stages and cortical cell types, with lowered expression in year 1-2 and in glial progenitors (**Figs. S6c, d**).

**Figure 6.**
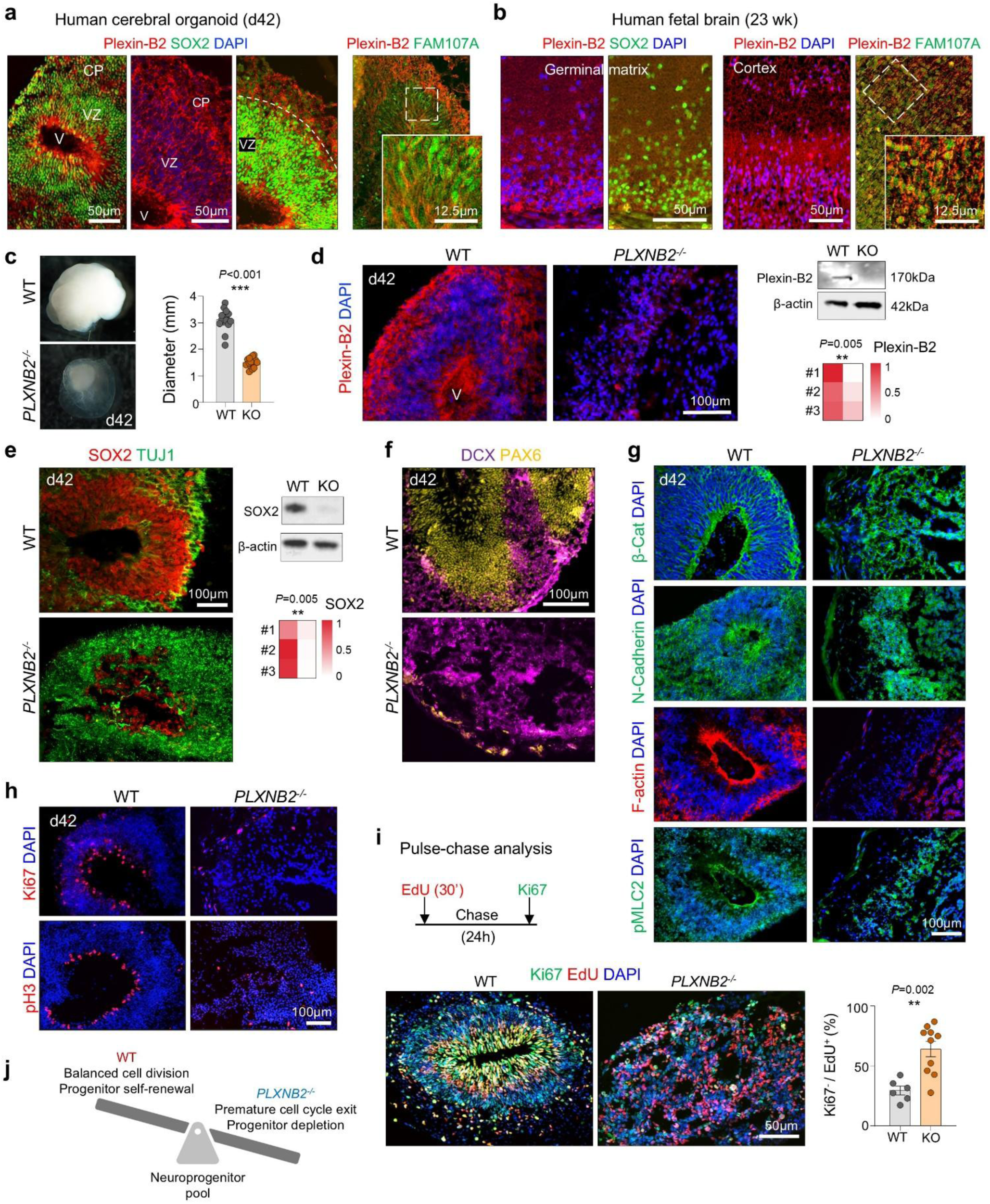
Plexin-B2 deficiency disrupts cortical development in human cerebral organoids. **a)** IF of day 42 cerebral organoids shows broad Plexin-B2 expression in the ventricular zone (VZ; SOX2⁺) and cortical plate (CP), including FAM107A⁺ outer radial glia. **b)** IF of human fetal brain at 23 gestational weeks reveals broad Plexin-B2 expression in the SOX2⁺ germinal matrix and developing cortex, including FAM107A⁺ cells. **c)** Representative images and quantification of organoid diameters show reduced size in *PLXNB2^⁻/⁻^* organoids (WT, n = 15; KO, n = 16). Bar graphs represent mean ± SEM; unpaired two-tailed Student’s t-test. **d)** Left, IF demonstrates loss of Plexin-B2 in d42 KO organoids. Right, Western blot confirms Plexin-B2 ablation, with β-actin as loading control. Quantification from n = 3 independent experiments. unpaired two-tailed Student’s *t*-test. **e)** Left, IF reveals disrupted architecture of KO organoids with shrinkage of SOX2^+^ VZ. Right, Western blot and heatmap show reduced SOX2 and increased TUJ1. n *=* 3 independent cultures per condition; unpaired two-tailed Student’s t-test. **f)** KO organoids exhibit expanded but disorganized DCX⁺ neuroblasts and reduced PAX6⁺ progenitors. **g)** Disrupted neuroepithelial organization in KO organoids, with diffuse β-catenin, N-cadherin, and reduced apical F-actin enrichment. **h)** WT organoids contain a dense apical ring of proliferating Ki67⁺/pH3⁺ cells, which was reduced and mislocalized in KO organoids. **i)** EdU pulse-chase assay design (30 min pulse, 24 h chase). KO organoids showed increased cell-cycle exit (i.e. fraction of Ki67⁻ cells among EdU⁺ cells). n *=* 6 fields from 3 independent organoids for each condition; unpaired two-tailed t-test. **j)** Model: Plexin-B2 maintains progenitor pool homeostasis by regulating the timing of neuronal differentiation. Loss of Plexin-B2 leads to premature cell-cycle exit, precocious neuronal differentiation, and progenitor depletion.

Compared to WT organoids, *PLXNB2*^-/-^ organoids were markedly smaller, lacked ventricular structures, and displayed irregular cystic cavities with disorganized SOX2⁺ or PAX6^+^ progenitors, but expansion of TUJ1^+^ and DCX^+^ populations (**Fig. 6c-f**). Deletion of Plexin-B2 protein was confirmed by IF and Western blot (**Fig. 6d**). Re-expression of CRISPR-resistant Plexin-B2 rescued ventricular morphology, confirming specificity (**Fig. S6a)**. Loss of Plexin-B2 also disrupted epithelial polarity: β-catenin, N-cadherin, and F-actin were mislocalized, and lamination markers (MAP2, TAU, vimentin, TBR1, CTIP2) revealed severe cortical disorganization (**Fig. 6g**; **Fig. S6e**).

To dissect the temporal requirement of Plexin-B2 in neurogenesis, we applied doxycycline (Dox)- induced shRNA knockdown (KD) (**Fig. S7a, b**). The KD studies demonstrated a critical temporal requirement: early (d14 or d21) *PLXNB2* depletion caused more severe ventricular collapse and FN disorganization than late KD (**Fig. S7a-d**).

Mechanistically, KO organoids exhibited reduced Ki67⁺ cycling progenitors, fewer apical pH3⁺ mitoses, and premature cell-cycle exit by EdU/Ki67 labeling (**Fig. 6h, i**). Early cell cycle exit was also observed in organoids with Pleixn-B2 KD (**Fig. S7e**). Thus, Plexin-B2 preserves neuroepithelial cohesion; its loss disrupts apical polarity, accelerates cell-cycle exit and neuronal differentiation, depletes neuroprogenitor pool, and ultimately impairs cortical morphogenesis (**Fig. 6j**).

### Neurogenic lineage instability and accelerated trajectories in Plexin-B2 KO organoids

To understand the molecular underpinnings of the defects observed in KO organoids, we performed snRNA-seq analysis on day 42 (d42) cerebral organoid (n=3 per genotype). This revealed distinct lineage states between WT and KO: WT organoids contained radial glia and immature/mature excitatory neurons, whereas KO organoids composed of subplate-like neurons (the earliest-born cortical neurons ^28^), maturing excitatory neurons, and also aberrant mesenchymal- or Schwann cell-like lineages (**Fig. 7a-d**). These data indicate an accelerated neurogenic trajectory with lineage switching and instability of neuronal identity in mutant organoids.

**Figure 7.**
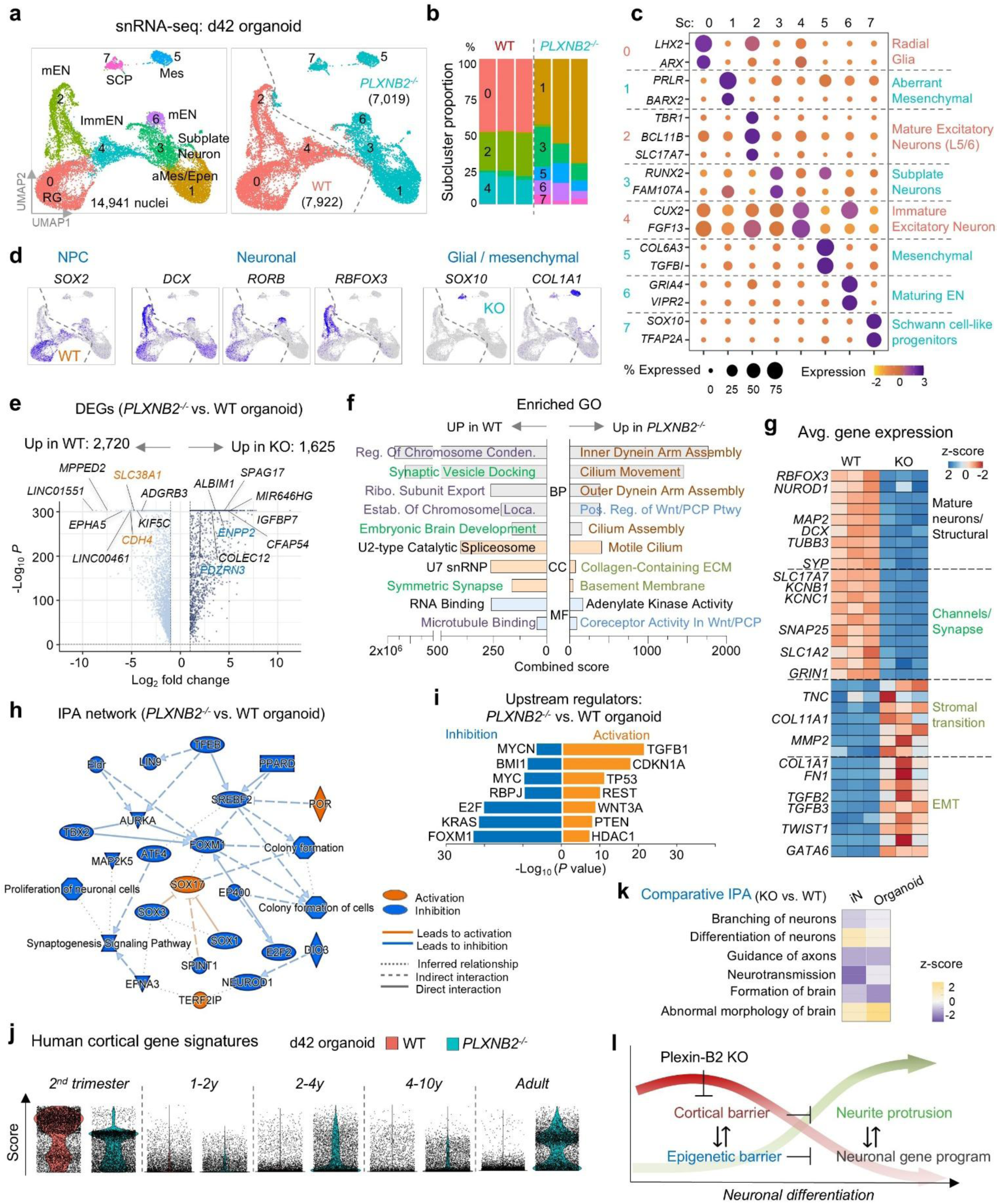
Accelerated neurogenic trajectory in *PLXNB2^-/-^* organoids is accompanied by lineage instability. **a)** UMAP embedding of snRNA-seq profiles from day 42 cerebral organoids (n = 3 replicates per genotype). EN, excitatory neurons; Mes, mesenchymal-like; Epen, ependymal-like; SCP, Schwann cell precursor/neural crest-like. b) Stacked bar plots show loss of RG (sc0) and expansion of subplate-like (sc3) and maturing EN (sc6) populations in *PLXNB2^⁻/⁻^* organoids, reflecting accelerated neurogenesis and lineage imbalance. c) Dot plot showing expression of marker genes across annotated subclusters. d) Feature plots highlighting distinctive marker gene expression in WT vs. *PLXNB2^⁻/⁻^* subclusters. e) Volcano plot of DEGs (adj. *P* < 0.05, log_2_FC > 1) with selected genes labeled. f) GO enrichment analysis of DEGs, grouped by biological process (BP), cellular component (CC), and molecular function (MF), color-coded by theme. g) Heatmap showing functional gene groups: WT organoids upregulated mature neuronal and synaptic genes, whereas *PLXNB2^⁻/⁻^* organoids upregulated stromal/EMT-associated genes, indicating lineage instability and aberrant mesenchymal-like states. h) IPA network analysis revealed broad suppression of progenitor/neuronal regulatory pathways in KO organoids, with limited activation of developmental branching and stress-response regulators. i) Predicted upstream regulators of DEGs, including suppression of proliferation- and neurogenesis-associated TFs and activation of senescence and stress regulators. j) Transcriptome scoring against human cortical developmental signatures (Velmeshev et al., 2023) with WT organoid cells primarily aligning with 2^nd^ trimester signatures, whereas *PLXNB2^⁻/⁻^* organoids also align with postnatal/adult signatures, indicating accelerated developmental age. k) Comparative IPA of *PLXNB2*^⁻/⁻^vs. WT in D36 iNs and d42 organoids revealed convergent alterations in neuronal differentiation and brain morphology pathways, supporting both models for Plexin-B2-linked neurodevelopmental defects. l) Model: Plexin-B2 enforces a cortical mechanical barrier integrated with an epigenetic barrier to safeguard the timing of neuronal differentiation. Loss of Plexin-B2 lowers this barrier, leading to premature epigenetic reprogramming, accelerated neurogenesis, and lineage instability.

DEGs highlighted ECM remodeling, ciliary pathways, and glial/mesenchymal programs (*FN1, COL11A1, SOX10, TGFB2*) and Wnt/PCP signaling in KO organoids, while WT upregulated neuronal maturation and synaptic genes (*MAP2, SYP, GRIN1, KCNB1*) (**Fig. 7e-g**).

IPA network and upstream regulator analyses indicated inhibition of progenitor maintenance factors (MYC, KRAS, FOXM1) and activation of SOX17 (a regulator of early lineage branching ^29^) and senescence/stress regulators (p21, TP53, PTEN), alongside epigenetic repressors (REST, HDAC1) (**Fig. 7h, i**). Developmental signature scoring using human cortical brain atlas further showed that KO organoid cells prematurely adopted late/postnatal cortical profiles, in contrast to WT cells that aligned with mid-gestational stages (**Fig. 7j**). Comparative analysis with iN cultures confirmed convergence of differentiation defects across 2D and 3D systems (**Fig. 7k**). Together, these data support Plexin-B2’s role in enforcing a cortical mechanical barrier which is coupled with an epigenetic barrier to regulate developmental timing and lineage commitment (**Fig. 7l**).

### Plexin-B2 function requires GAP activity and mechanosensitive extracellular domain

To elucidate Plexin-B2 signaling pathway during neuronal differentiation, we conducted structure-function analysis developed previously ^11,30^ (**Fig. S8a**). Mutational analysis pinpointed the Ras-GAP domain and the mechanosensitive extracellular ring as critical. Specifically, in rescue experiments, GAP-domain (mGAP) or extracellular truncation (ΔECTO) mutants failed to rescue the KO phenotypes in both iN and organoid assays, in contrast to mutants in Rho-binding domain (mRBD) or deletion of the C-terminal PDZ-binding motif (ΔVTDL) that reverted the KO phenotypes (**Fig. S8b, c**).

Next, we examined locked ring (LR) mutations of Plexin-B2 in which the extracellular ring is disulfide-bridged that render Plexin-B2 insensitive to mechanical stimuli, but semaphorin-responsive ^30,31^. LR mutants also displayed accelerated differentiation in neural induction assays, similar to *PLXNB2* KO (**Fig. S8d**). In 3D assays, LR mutant organoids were smaller than WT, with disorganized SOX2^+^ or PAX6^+^ progenitor cells intermingled with TUJ1^+^ and DCX^+^ populations, mirroring *PLXNB2*^-/-^ phenotypes albeit with less severity (**Fig. S8e-f**). Thus, Plexin-B2 requires both intracellular GAP activity and extracellular mechanical flexibility to function as a cortical checkpoint for neuronal differentiation.

## DISCUSSION

Neuronal differentiation has traditionally been understood as the outcome of gene regulatory cascades and transcriptional programs. Our study reveals a new principle in that neuronal fate commitment is also gated by a cortical mechanical barrier that integrates with nuclear epigenetic checkpoints to ensure temporal fidelity of neurogenesis.

We show that cortical actin tension, regulated by Plexin-B2, functions as a mechanical barrier that prevents premature neurite protrusion. Lowering this barrier, through Plexin-B2 deletion, function-blocking anti-Plexin-B2 nanobodies, or pharmacological actin destabilization, was sufficient to accelerate neurite initiation and advance morphodynamic transitions. This finding aligns with earlier work showing that low doses of actin polymerization inhibitors enhance neurite outgrowth by suppressing actomyosin contractility ^32^. Similarly, soft matrix substrate also promotes neuronal differentiation of hESCs ^33^. Mathematical simulations further illustrated that effective morphogenesis requires an optimal balance between membrane tension and membrane-cytoskeletal energy. Transcriptomic profiling highlighted a central role for Plexin-B2 in controlling cell mechanics and morphological transformation during neuronal differentiation. Finally, structure-function analysis revealed that both Plexin-B2’s Ras-GAP activity and its flexible extracellular ring domain are indispensable for the checkpoint function, establishing Plexin-B2 as a mechano-gatekeeper of neuronal morphogenesis.

Beyond morphogenesis, Plexin-B2 deficiency also resulted in precocious neuronal gene expression, indicating a connection of mechanodynamics and nuclear gene programs. In line with this, Plexin-B2 deletion precipitated accelerated epigenetic reprogramming, with early upregulation of lamin A/C and Tet3, recapitulating a state normally seen at later differentiation. This convergence of cytoskeletal mechanics and nuclear regulation reveals that cortical actin and epigenetic barriers act in concert to safeguard developmental timing. Although the precise mechanisms synchronizing morphodynamics with nuclear gene programs remain to be determined, earlier research has shown that mechanical cues can activate mechanosensitive pathways and alter nuclear envelope properties ^34^. Our findings thus expand the framework of corticogenesis from gene-centric to mechano-epigenetic models of fate control.

In human cerebral organoids and fetal brain, Plexin-B2 was enriched in ventricular zones and outer radial glia, consistent with a role in progenitor maintenance. The transient decline of Plexin-B2 during neurogenesis suggests a gating function, preventing premature differentiation until appropriate developmental cues converge. Loss of Plexin-B2 disrupted apical polarity of neuroepithelium and depleted the progenitor pool through premature cell-cycle exit, resulting in smaller organoids with cyst-like structures and lineage instability. These defects parallel neural tube closure phenotypes in *Plxnb2* knockout mice and align with recent reports of rare pathogenic *PLXNB2* variants in patients with intellectual disability, underscoring clinical relevance. Hence, by extending Plexin signaling beyond its classic role in axon guidance to biomechanical regulation, our study provides a new framework to understand neurodevelopmental disorders through the lens of mechanomorphogenesis.

While prior efforts for optimizing induced neurons focused on stepwise application of differentiation factors ^15,35^, we show here that lowering the cortical barrier halved the timeline for induced neuron derivation (from ∼36 to <18 days). *PLXNB2^-/-^* iNs not only displayed stereotypical neuronal morphology with long processes and dense neurite networks, they also upregulated neuronal markers, downregulated progenitor genes, and acquired more mature transcriptomic and electrophysiological states with higher firing rates and enhanced network synchrony after further maturation in co-cultures with astrocytes. Importantly, temporal alignment of external cues with intrinsic morphodynamic states stabilized neuronal identity, offering a strategy to overcome lineage instability. Nanobody-mediated Plexin-B2 inhibition further provides a tractable tool to temporally tune neuronal differentiation. These findings suggest new approaches for accelerating the generation of functionally mature neurons for CNS disease modeling and regenerative medicine.

In conclusion, our work establishes cortical actin tension as a mechanical barrier to premature neuronal commitment. By integrating a cortical mechanical checkpoint with an epigenetic nuclear barrier, our study advances a unifying framework for neuronal differentiation. This model reshapes our understanding of corticogenesis and also provides insight into neurodevelopmental disorders and opens translational opportunities to accelerate neuronal maturation to generate induced neurons for CNS disease modeling and cell therapy.

## METHODS

### hESCs

H9 human embryonic stem cells (hESCs) ^36^ were obtained from the University of Wisconsin and maintained on Matrigel (#354230, Corning)-coated culture dishes in mTeSR1 medium (#100-0274, Stemcell Technologies). Medium was replaced every other day. All hESC experiments were approved by the Institutional Review Board and Ethics Committee at the Icahn School of Medicine at Mount Sinai.

### Neural progenitor differentiation

hESCs were differentiated into NPCs using the STEMdiff SMADi Neural Induction system (#08581, Stemcell Technologies). Briefly, hESCs were maintained on a Matrigel-coated 6-well plate in mTeSR Plus Basal Medium (#100-0274, Stemcell Technologies) before neural induction. At appropriate stage, cells were dissociated using Accutase Cell Detachment Solution (#SCR005, MilliporeSigma); 2×10^6^ cells (2x10^5^ cells/cm^2^) were transferred to a new Matrigel-coated 6-well plate in 2 ml STEMdiff Neural Induction Medium supplemented with SMADi (#08580, STEMCELL Technologies) and 10 μM Y-27632 (#72302, Stemcell Technologies). The medium (without Y-27632) was replenished daily with warmed (37 °C) STEMdiff Neural Induction Medium + SMADi until completion of the neural induction protocol. Cells were passaged three times (days 9, 15, and 21) and reseeded onto a new Matrigel-coated 6-well plate at 1.8 × 10^6^ per well (125,000 cells/cm^2^). Each transfer was supplemented with 10 μM Y-27632 (ROCK inhibitor) on day 1 of seeding to increase cell survival.

### Neuronal maturation

Differentiation of forebrain and midbrain neuronal precursors into mature neurons was performed according to the manufacturer’s instructions using the STEMdiff Forebrain Neuron Differentiation Kit (08600#, Stemcell Technologies) for forebrain-type neurons, and the STEMdiff Midbrain Neuron Differentiation Kit (#100-0038, Stemcell Technologies) for midbrain neurons. Following the STEMdiff SMADi Neural Induction system as described above, on day 22 of the protocol (following passage 3), neural induction medium was aspirated, and STEMdiff Forebrain or Midbrain Neuron Differentiation Medium was introduced. A daily medium change with warm (37 °C) differentiation medium was performed until cells reached 80-90% confluence (days 27-28). Cells were then passaged and reseeded onto a new Matrigel-coated 6-well plate at 5.4 × 10^5^ per well (6 × 10^4^ cells/cm^2^) in STEMdiff Forebrain (#08605, Stemcell Technologies) or Midbrain (#100-0041) Neuron Maturation Medium, accordingly. Cells were incubated at 37 °C, with change of half the medium every 2 - 3 days. After 8 days in maturation medium, cells were analyzed for neuronal phenotypes.

### Molecular dynamics simulations

The molecular dynamics (MD) simulations were based on a coarse-grained cell models as described in previous publications ^11,30^. Briefly, molecular dynamics modelling was performed using a cell model composed of beads representing the cell membrane, the nuclear membrane, actin filaments, and heads of actin filaments. The nuclear membrane is connected to the cell membrane through actin filaments. Connections between the beads are modelled by springs of elastic constant κ. The total number of beads per cell in the model was n = 700. Additional details on MD calculations are described in **Supplementary File 1**.

### CRISPR genome editing for *PLXNB2* deletion

Plexin-B2 deletion in human cell lines by lentiviral CRISPR-Cas9 vectors has been described in a previous study ^11^. In brief, low passage hESCs were stably transduced with lentiviral particles produced in HEK293T cells using plenti-CRISPRv2 vectors encoding Cas9 and an sgRNA targeting either exon 2 of *PLXNB2* (sequence: GTTCTCGGCGGCGACCGTCA; Addgene #86152) or the EGFP coding sequence (sequence: GGGCGAGGAGCTGTTCACCG; Addgene #86153). Transduced cells were selected with puromycin (1 µg/ml, 7 days) and Western blot assay was used to confirm loss of the mature Plexin-B2 protein.

### Drug treatment during neuronal induction

WT and *PLXNB2^⁻/⁻^* hESCs were seeded at a density of 2 × 10^6^ cells (2 × 10^5^ cells/cm²) in Matrigel-coated 6-well plates with 2 ml STEMdiff Neural Induction Medium supplemented with dual-SMAD inhibition (SMADi) as described above. Beginning on day 0, cells were treated with either latrunculin A (F-actin inhibitor; #10010630, 0.5 μM; Cayman Chemical), jasplakinolide (F-actin stabilizer; #11705, 0.5 μM; Cayman Chemical), or vehicle control (DMSO; #D1281, Fisher Scientific). Drug-containing media were freshly prepared and replaced daily. Cultures were maintained under these conditions for 9 days, followed by passaging on day 9 and immunostaining on day 10.

### Live cell imaging

WT and *PLXNB2^⁻/⁻^* hESCs were seeded on day 0 at a density of 2 × 10^6^ cells (2 × 10^5^ cells/cm^2^) in Matrigel-coated 6-well plates with 2 ml STEMdiff Neural Induction Medium supplemented with dual-SMAD inhibition (SMADi). Cultures were maintained for 6 days with daily medium replacement using pre-warmed (37 °C) STEMdiff Neural Induction Medium + SMADi. Y-27632 was included only during the first 24 h after seeding. On day 6, cells were labeled with SPY555-Actin (#CY-SC202, 1:1000, Cytoskeleton), SPY650-Tubulin (#CY-SC203, 1:1000, Cytoskeleton), and NucSpot Live 488 (#40082, 1:1000, Biotium). Live-cell time-lapse imaging was carried out for 30 hrs on a Zeiss LSM 780 confocal microscope equipped with environmental chamber (37 °C, 5% CO_2_), with frames acquired every 20 min. After imaging, cells were fixed for endpoint analysis. Image processing and quantification were performed using Imaris software (Oxford Instruments)

### Generation of anti-Plexin-B2 nanobodies

Nanobodies were generated by immunizing a llama and an alpaca with the extracellular domain of human Plexin-B2 (recombinant protein, expressed as a fusion with llama Fc in Expi293 cells). Peripheral blood mononuclear cells (PBMCs) were isolated from the immunized animals and used to generate cDNA libraries of the variable domains of the heavy chains in heavy-chain-only antibodies (VHH). Three rounds of phage display were applied to enrich for VHH clones binding human Plexin-B2 with high affinity. Twenty-one selected VHH clones were expressed as VHH-human Fc nanobody proteins by Chinese hamster ovary (CHO) cells and assessed for binding properties and function-blocking activity in cell culture assays. For the experiments in this study, the function-blocking anti-Plexin-B2 VHH-hFc clones YR-25 and YR-38 were used. The VHH-hFc nanobody YR-43, raised against the unrelated human intracellular protein LRRK2, was used as a negative control.

### Neonatal astrocyte preparation for co-culture assay

For co-culture experiments, cortical astrocytes were isolated from neonatal (P1-3) mouse pups. Following removal of meninges, cortices were dissected into small pieces, washed in HBSS, and enzymatically dissociated using the Neural Tissue Dissociation Kit-T (#130-094-802, Miltenyi Biotec). Dissociated cells were plated on tissue culture dishes and maintained for 3-5 days in DMEM/GlutaMAX supplemented with 10% FBS and 1:100 penicillin-streptomycin (#15140122, Gibco). Astrocyte enrichment was subsequently performed using the anti-GLAST (ACSA-1) MicroBead Kit (#130-095-825, Miltenyi Biotec).

### Multielectrode array (MEA) recordings

Neuronal network activity was assessed using the Maestro MEA platform (Axion Biosystems). Cells were plated on a CytoView MEA 24 plate with 16 electrodes per well (#M384-tMEA-24W, Axion Biosystems). Induced neurons from WT hESCs differentiated for 36 days with standard protocol or from *PLXNB2^-/-^* hESCs generated by the accelerated 18 day-protocol were dissociated and replated onto MEA wells at a density of 60,000 cells per well (corresponding to ∼2-3 × 10^5^ cells/cm^2^). To support neuronal survival and maturation, primary mouse cortical astrocytes isolated from P1-3 pups (see astrocyte preparation above) were seeded onto MEA wells three days prior to neuronal transfer at a neuron-to-astrocyte ratio of 3:1 (∼20,000 astrocytes per well). Astrocytes were initially maintained in DMEM/GlutaMAX supplemented with 10% FBS, and after neuronal seeding, the medium was switched to forebrain neuronal maturation medium. Both WT and KO neurons were maintained under the same forebrain maturation protocol, and only forebrain-lineage neurons were used for MEA assays.

Cultures were maintained with half-medium replacement every 2 days. Electrical recordings were performed weekly for 3 weeks. Data acquisition and spike detection were carried out using AxIS Navigator software (v3.7.2, Axion Biosystems). Parameters analyzed included mean firing rate (Hz), synchrony index, and spike rate (spikes/s).

### Single-nucleus RNA-seq library preparation and sequencing

Single-nucleus suspensions were prepared from WT and *PLXNB2^-/-^*hESC-derived cultures collected at multiple time points during the differentiation protocol, as described in the Results section. Nuclei were isolated using a gentle lysis protocol optimized to preserve nuclear structure and RNA integrity (#130-128-024, Miltenyi Biotec). Isolated nuclei were counted, quality-checked via DAPI staining, and processed for library preparation using the Evercode Whole Transcriptome v2 kit (Parse Biosciences, Version 2.3-UM0022), following the manufacturer’s instructions. Final libraries were sequenced on an Illumina NovaSeq X platform using a 25B flow cell (300-cycle, standard), generating paired-end 150 bp reads. Libraries were sequenced to a depth of ∼50,000 reads per nucleus. Raw reads were processed with Evercode Whole Transcriptome Analysis software (Parse Biosciences), which performs barcode demultiplexing, alignment to the GRCh38 reference genome, and UMI-based quantification, generating gene-by-cell count matrices for downstream analysis.

### Single-nucleus RNA-seq data analysis

Gene-by-cell count matrices were imported into R and processed using the Seurat (v5.0.1) package ^37^. Quality control filtering retained nuclei with >2,000 detected genes, >1,000 UMIs, and <25% mitochondrial gene content. Across all samples, 44,085 nuclei passed QC and were retained for downstream analysis. WT and *PLXNB2^⁻/⁻^* datasets were merged into a single Seurat object for joint analysis to allow direct comparison across genotypes. Data were normalized, scaled, and the top 2,000 highly variable genes were selected for principal component analysis (PCA). Dimensionality reduction was performed using UMAP based on the first 10 principal components. Graph-based clustering was performed with the Seurat FindNeighbors function (using the first 10 PCs) followed by FindClusters at a resolution of 0.1.

Clusters were annotated based on canonical marker expression, the top 100 marker genes per cluster, and label transfer from a published human cortical atlas ^7^ using the Seurat FindTransferAnchors, TranferData, and addMetaData functions. Clusters co-expressing incompatible lineage markers were flagged as doublets and excluded from downstream analysis.

Differential gene expression analysis was performed in Seurat using the FindMarkers function (Wilcoxon rank-sum test). For functional enrichment, stringent thresholds were applied (adjusted *P* < 0.05 and |log₂ fold change| > 1, min.pct = 0.1) to define significantly up- or downregulated genes. In contrast, for Ingenuity Pathway Analysis (IPA, QIAGEN), a permissive cutoff (logfc.threshold = 0, min.pct = 0) was used to provide the full ranked gene list required for pathway-level inference. Gene Ontology (GO) enrichment analyses were conducted separately for upregulated and downregulated genes using the ENRICHR platform (https://maayanlab.cloud/Enrichr/) ^38^. Gene sets that were analyzed were GO Biological Process 2025, Cellular Component 2025, and Molecular Function 2025. Additional pathway enrichment was performed with WikiPathways (Human, 2024 release) and KEGG (2025 release) gene sets.

### Data visualization

Bubble plots were generated in ggplot2 (v3.4.4) from clusterProfiler enrichment results, with circle size representing gene counts and color mapped to -log_10_(*p.adjust*). Heatmaps were generated in R using pheatmap (v1.0.12), with row-wise z-score scaling applied to average expression values per condition. Dot plots for cluster annotation were generated using the Seurat DotPlot function, with further customization in ggplot2. In dot plots, dot size represents the percentage of nuclei expressing each gene and color reflects scaled average expression values.

### Cerebral organoid derivation from hESCs

hESC-derived cerebral organoids were generated using the STEMdiff Cerebral Organoid Kit (#08570, Stemcell Technologies). In brief, WT and *PLXNB2^-/-^*hESCs (<30 passages) were dissociated with Accutase and seeded at 9,000 cells/well into 96-well ultra-low attachment (ULA) U-bottom plates (#7007, Corning) to form embryoid bodies. On day 5, aggregates were transferred to 24-well ultra-low attachment plates (#3473, Corning) for neural induction (∼48 h). On day 7, organoids were embedded in Matrigel (#354230, Corning) and moved to 6-well ULA plates for expansion. From day 10, cultures were placed on an orbital shaker (65 rpm for 6-well) in maturation medium, with medium changes every 2-3 days, and maintained until day 42. For snRNA-seq, organoids were processed using Cultrex Organoid Harvesting Solution (#3700-100-01, R&D).

### Doxycycline-inducible Plexin-B2 knockdown

Inducible knockdown (KD) of *PLXNB2* was generated as previously described ^11^. Briefly, hESCs were transduced with Tet-pLKO-Puro-based lentiviral vectors expressing doxycycline (Dox)-inducible shRNAs against *PLXNB2* (pLKO-Tet-On-PLXNB2-shRNA1 or -shRNA2; backbone Addgene #21915, shRNA1 Addgene #98399, shRNA2 Addgene #98400) or a non-targeting control (pLKO-Tet-On-shRNA-Ctrl). Stable lines were established by puromycin selection (1 µg/ml, 7d; #P8833, Sigma-Aldrich). Knockdown was induced with 1 µg/ml doxycycline (#D5207, Sigma-Aldrich) added to the culture medium and replenished at each medium change. These hESC lines (*PLXNB2* shRNA and control) were then used to generate cerebral organoids following the STEMdiff protocol described above. For temporal induction experiments, Dox administration was initiated at different stages of organoid development (day 14, 21, 28, or 35) and maintained until analysis (day 42); parallel no-Dox controls were processed in the same manner. KD efficacy was assessed by immunofluorescence and Western blot.

### Lentiviral rescue and signaling mutants

Rescue experiments with Plexin-B2 mutant forms were performed as described previously ^11^. Briefly, a CRISPR-resistant *PLXNB2* cDNA was generated by site-directed mutagenesis and cloned into the pLenti-PGK vector (Addgene #19067) to produce the rescue construct (pLV-*PLXNB2*). Signaling mutants were derived from this backbone, including mGAP (R1391A/R1392G), mRBD (LSK1558-1560→GGA), ΔVTDL (C-terminal PDZ-binding motif deletion), and ΔECTO (extracellular deletion, aa 30-1189). Lentiviral particles were produced in HEK293T cells and used to transduce *PLXNB2^-/-^* hESCs, with puromycin selection to establish stable lines for organoid assays.

### Pulse-chase proliferation assay

Organoids were subjected to a pulse-chase labeling protocol to assess progenitor cell cycle status. Briefly, day-42 cerebral organoids were incubated with 10 µM EdU for 30 min (pulse) and then washed and returned to fresh maturation medium for a 24 h chase period. Organoids were fixed in 4% paraformaldehyde, cryoprotected, embedded in OCT, and sectioned at 15 µm thickness. Incorporated EdU was detected using the Click-iT EdU Cell Proliferation Kit for Imaging with Alexa Fluor 594 dye

(#C10339, Thermo Fisher). Following EdU detection, cryosections were immunostained with anti-Ki67 antibody to label proliferating nuclei, and counterstained with DAPI. The proportion of cells having undergone cell cycle exit was quantified as fraction of Ki67^-^ EdU⁺ cells relative to total EdU⁺ cells.

### Antibodies

#### Primary

anti-β-catenin (host: mouse, BD Bioscience 610153, 1:200)

anti-CTIP2 (host: rat, Abcam ab18465, 1:250)

anti-DCX (host: guinea pig, EMD Millipore AB2253, 1:500)

anti-FAM107A (host: rabbit, Atlas antibodies 055888, 1:200)

anti-FN (host: rabbit, EMD Millipore ab2033, 1:200)

Anti-FOXG1(host: rabbit, Invitrogen 702554, 1:250)

anti-GFAP (host: chicken, Aves Labs GFAP, 1:500)

anti-Ki67 (host: rabbit, Abcam 16667, 1:300)

anti-LaminA/C (host: mouse, Invitrogen mab636, 1:200)

anti-N-cadherin (host: mouse, BD Bioscience 610920, 1:200)

anti-NeuN (host: chicken, Aves Lab NUN, 1:300)

anti-NF-H (host: chicken, EMD Millipore AB5539, 1:1,000)

anti-OCT4 (host: mouse, Abcam ab184665, 1:500)

anti-MAP2 (host: chicken, Aves Lab MAP, 1:1000)

anti-MECP2 (host: mouse, Biolegend 814001, 1:300)

anti-Nanog (host: rabbit, Abcam ab109250, 1:200)

anti-Nestin (host: mouse, R&D systems IC1259G, 1:350)

anti-PAX6 (host: mouse, Abcam ab78545, 1:200)

anti-pH3 (host: rabbit, Cell Signaling 9701, 1:200)

anti-Plexin-B2 (extracellular domain) (host: sheep, R&D systems AF5329, 1:300)

anti-pMLC2 (host: rabbit, Cell Signaling 3671, 1:200)

anti-Sema4B (host: rabbit, Proteintech 16934-1-AP, 1:300)

anti-Sema4C (host: sheep, R&D systems AF6120, 1:300)

anti-Sox2 (host: rabbit, Abcam ab97959, 1:200)

anti-Tau (host: mouse, Invitrogen AHB0042, 1:200)

anti-TBR1 (host: rabbit, Abcam 31940, 1:300)

anti-TET3 (host: rabbit, Abcam 139311, 1:300)

anti-Tuj1 (host: mouse, Biolegend 801202, 1:1,000)

anti-Vimentin (host: chicken, Novus Bio NB300-223, 1:350)

For F-actin staining: Phalloidin-Alexa594 (Invitrogen A12381, 1:400)

#### Secondary

Alexa Fluor 488, 594, or 647-conjugated donkey anti-goat, -rabbit, -rat, -mouse, and -guinea pig IgG (Jackson ImmunoResearch Laboratories, 1:300).

#### Western blot

anti-β-actin (host: rabbit, Sigma A1978, 1:10,000),

anti-Plexin-B2 (ecd) (host: sheep, R&D Systems AF5329, 1:500)

anti-SOX2 (host: rabbit, Abcam ab97959, 1:10,000)

### Immunocytochemistry and immunofluorescence

For immunocytochemistry (ICC) of cultured cells, cells were fixed with 3.7% formaldehyde in PBS at room temperature for 10 min, washed three times with PBS, and incubated in blocking buffer (5% donkey serum and 0.3% Triton X-100 in PBS) for 1 h. Primary antibodies were applied in dilution buffer (PBS with 1% BSA and 0.3% Triton X-100) and incubated overnight at 4 °C. Secondary antibodies were added in dilution buffer at room temperature for 1 h together with nuclear counterstaining using DAPI (1:1,000; Thermo Fisher).

For immunofluorescence analysis of cerebral organoids, day 42 organoids were fixed in 4% paraformaldehyde in PBS at 4 °C for 15 min, washed three times in PBS, cryoprotected sequentially in 12.5% sucrose (overnight, 4 °C) and 25% sucrose (overnight, 4 °C), then embedded in O.C.T. compound (#4585, Fisher HealthCare) and frozen on dry ice. Organoids were sectioned at 15 μm thickness using a cryostat (Leica), and cryosections were processed for antibody staining following the ICC protocol described above.

### Western blotting

For Western blotting, cells were lysed with RIPA buffer (Sigma) containing protease and phosphatase inhibitors. Protein concentrations were determined using a BCA assay (Thermo Scientific). Proteins were resolved by SDS-PAGE on 4-12% polyacrylamide NuPAGE gels (Invitrogen) and transferred onto nitrocellulose or PVDF membranes (Li-Cor Biosciences) with the XCell transfer system (Invitrogen). Membranes were incubated at 4 °C overnight with primary antibodies and then for 1 h with secondary donkey antibodies coupled to IRDye 680 or 800 (Li-Cor). The fluorescent bands were detected with an Odyssey infrared imaging system (Li-Cor Biosciences).

### Statistical analyses

We used unpaired two-tailed Student’s *t*-tests for comparisons between two groups. For comparisons involving multiple groups, we applied two-way ANOVA with Dunnett’s multiple comparisons test. For multiple field analyses, nested structures for both the t-test and the *one*-way ANOVA were used. All statistical analyses were conducted using GraphPad Prism version 10, using the NEJM (New England Journal of Medicine) style setting for reporting of *P* values. Statistical significance was defined as follows: * *P* ≤ 0.01, \*\**P* ≤ 0.01, \*\*\**P* ≤ 0.001.

## Acknowledgements

We thank Theodore Hannah, Icahn School of Medicine at Mount Sinai, for help with organoid experiments and quantifications, and Zou and Friedel lab members for constructive comments throughout the project.

## Author contributions

D.H., C.J.A., R.H.F., and H.Z. conceived and designed the study. D.H., C.J.A., and J.L. performed neural induction and organoid culture studies, as well as single cell sequencing. M.E. and L.S. conducted the bioinformatic analysis of single-cell transcriptomic data. S.D., H.T., S.K., X.L., and Z.Y. generated nanobodies. N.M.T. supported studies with human fetal sections. Y.P. and S.I.R. conducted analyses of human transcriptomic data. M.R.L., R.A.D., G.O.R., and J.P.R.F.M. conduction molecular dynamics simulations. Manuscript and figures were prepared by D.H., C.J.A., R.H.F., and H.Z.

## Data availability

The RNA-Seq data has been deposited at the NCBI Gene Expression Omnibus (GEO) database under accession number GSE XXXXXX.

