## Supplementary material for "Cortical tension as a mechanical barrier to safeguard against premature differentiation during neurogenesis": Supplemetary Materials

Halperin et al.

**Supplemental Movie S1**

Live-cell imaging over 42 hours of WT and PB2 KO D6 cells labeled with NucSpot, SPY-tubulin, and SPY-actin.

**Supplemental Figures S1 - S8.**

See pages below.

**Supplemental Table S1.**

Overview of cell numbers in single nucleus RNA sequencing samples after quality filtering.

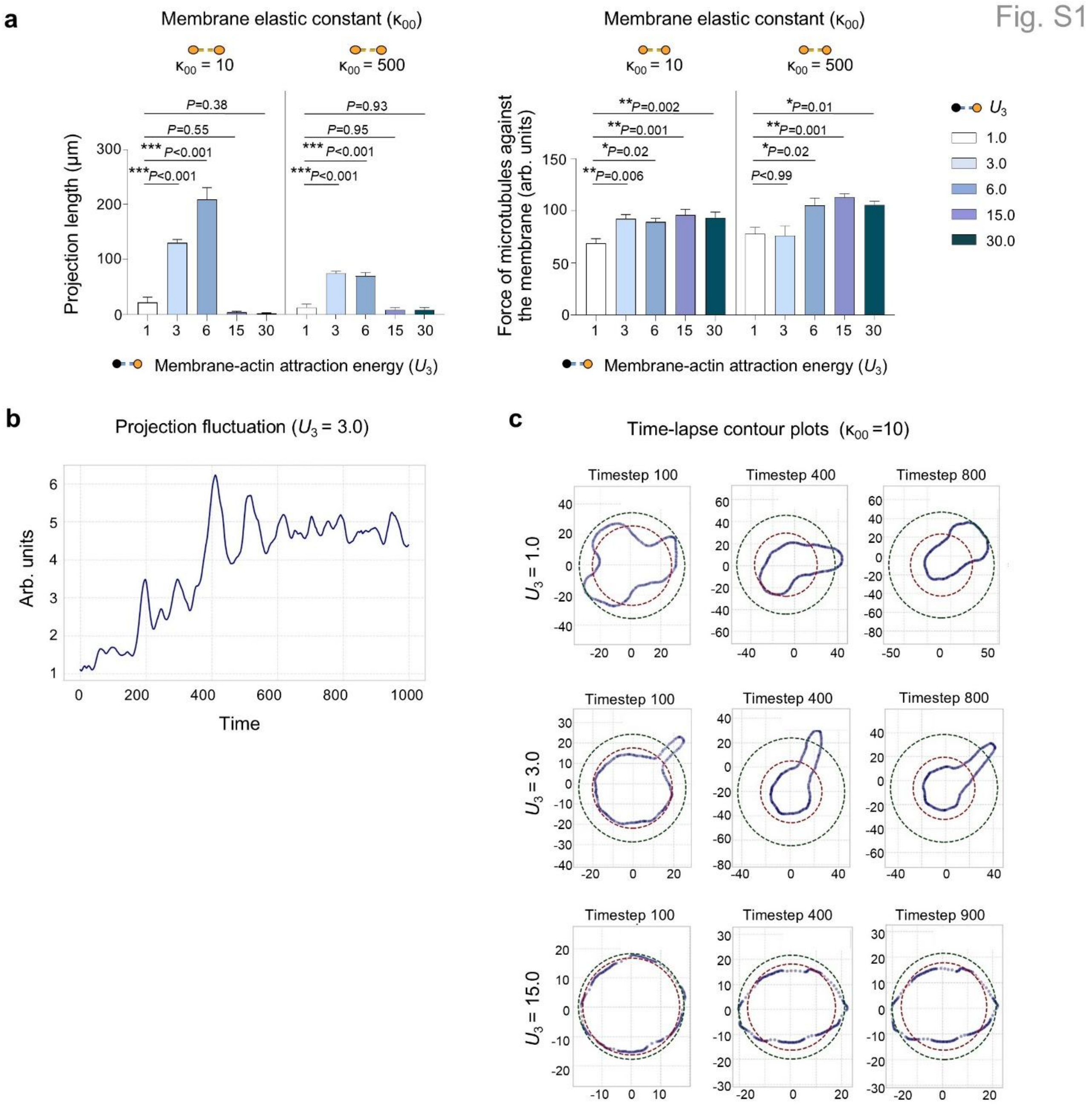

**Figure S1. Molecular dynamics modeling of cortical actin tension during neurodifferentiation.**

**a)** Simulated neurite projection area (left) and microtubule pushing force (right) under two membrane elastic constants ($\kappa_{00}$= 10 vs. 500) and varying membrane-cytoskeleton attraction energies ($U_3$). At low membrane tension ($\kappa_{00}$= 10), projection area increases markedly at moderate $U_3$ values, indicating that soft membranes favor neurite protrusion. Extremely low $U_3$ values impaired projection stability, reflecting insufficient actin-membrane coupling. At high membrane tension ($\kappa_{00}$ = 500), projections were suppressed across all $U_3$ conditions. Microtubule pushing force remained largely unchanged.

Bar graphs (for projection length and force against membrane) represent mean ± SEM, one-way ANOVA with Dunnett's multiple-comparison test.

**b)** Time-course analysis of projection length at moderate $U_3$ (3.0) with low membrane tension ($\kappa_{00} = 10$) shows dynamic fluctuations of extensions, consistent with active and flexible membrane states that promote neurite protrusion.

**c)** Simulated cell shape dynamics under varying $U_3$ values with constant membrane tension ($\kappa_{00} = 10$). Contour plots show evolving cell boundaries (blue) at selected time points, with dashed red (nuclear center) and green (cellular boundary) reference circles. At $U_3 = 3.0$, cells formed a stable polarized projection with nuclear compaction and displacement, mimicking early neurite emergence. Low $U_3$ (1.0) yielded unstable morphologies without clear axis formation, while high $U_3$ (15.0) constrained cells to near-spherical shapes with minimal protrusion. These results suggest that balanced actin-membrane coupling and cortical tension are critical for stable, directed neurite initiation.

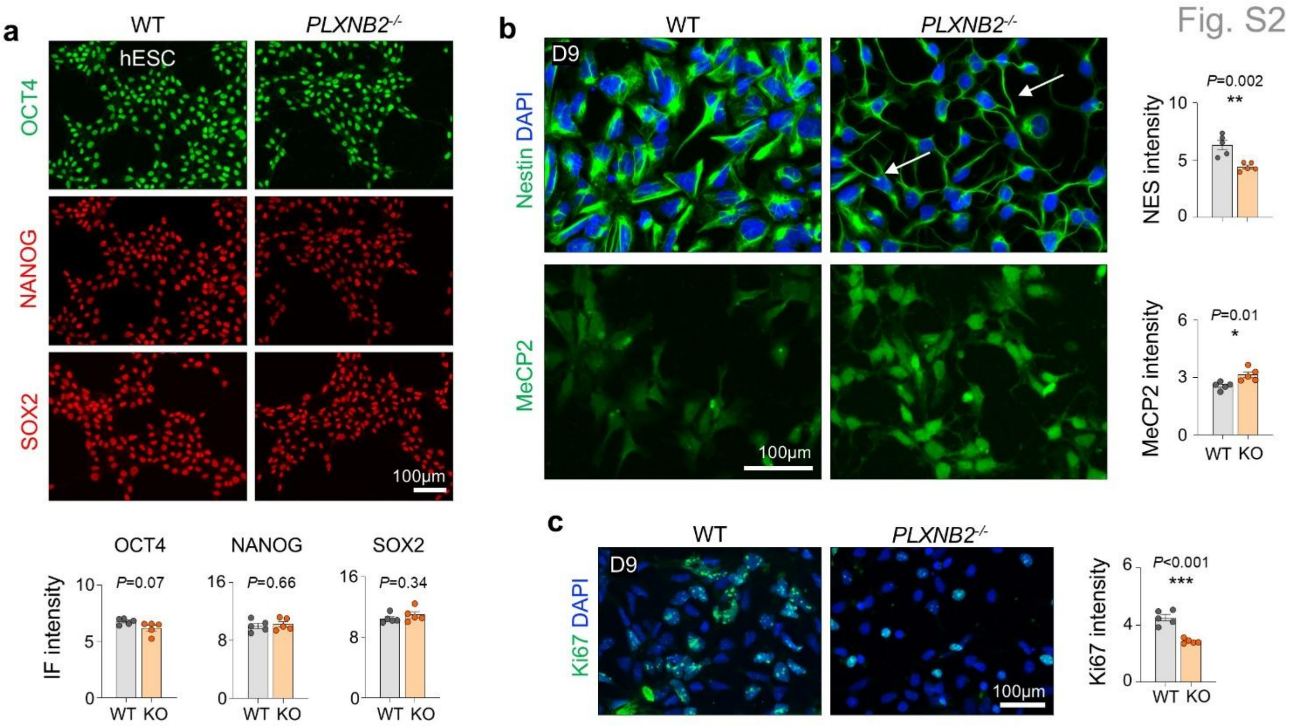

**Figure S2. Plexin-B2 KO cells display early neuronal lineage commitment.**

**a)** IF demonstrates comparable expression of pluripotency markers in WT and *PLXNB2*$^{-/-}$ hESCs.

**b)** IF shows reduced Nestin and increased MeCP2 expression in D9 *PLXNB2*$^{-/-}$ cells relative to WT. KO cells also displayed morphological changes with elongated protrusions (arrows), a sign of premature neuronal differentiation.

**c**) IF for proliferation marker Ki67 reveals reduced proliferative activity in D9 *PLXNB2*$^{-/-}$ cultures. Quantification based on n = 5 fields (each averaged from ~50 cells per field) for each condition, two-tailed nested t-test. Bar graphs represent mean ± SEM.

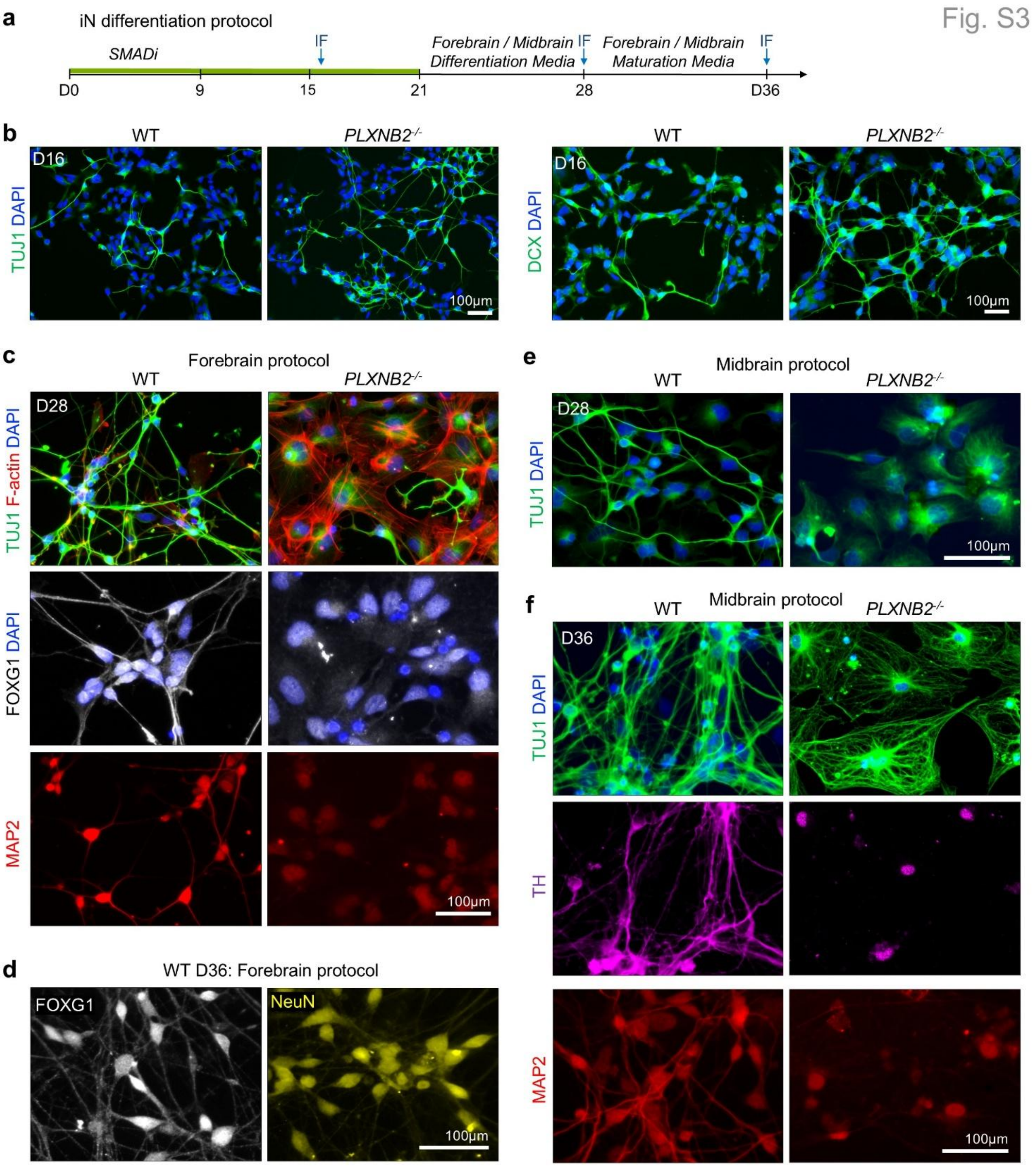

**Figure S3. Lineage instability in *PLXNB2*$^{-/-}$ cells under mismatched media conditions.**

**a)** Timeline of forebrain or midbrain differentiation protocols for iNs.

**b)** IF at D16 of iN protocol shows denser networks of DCX$^{+}$ or TUJ1$^{+}$ neurite projections in *PLXNB2*$^{-/-}$ cells as compared with WT cells.

**c**) At D28 of forebrain differentiation protocol, *PLXNB2*$^{-/-}$ cells display aberrant morphology and loss of stable neuronal marker gene expression, in contrast to WT cells that displayed organized TUJ1$^{+}$ and MAP2$^{+}$ neuronal networks and expression of forebrain marker FOXG1.

**d)** WT D36 iNs from the forebrain protocol show expression of NeuN and forebrain marker FOXG1.

**e)** At D28 of midbrain differentiation protocol, WT cells maintained organized TUJ1$^{+}$ neuronal networks, but *PLXNB2*$^{-/-}$ cells have acquired an aberrant morphology.

**f)** WT D36 cells treated with midbrain maturation media. *PLXNB2*$^{-/-}$ cells show loss of organized TUJ1 and MAP2 networks and lack expression of midbrain marker tyrosine hydroxylase (TH).

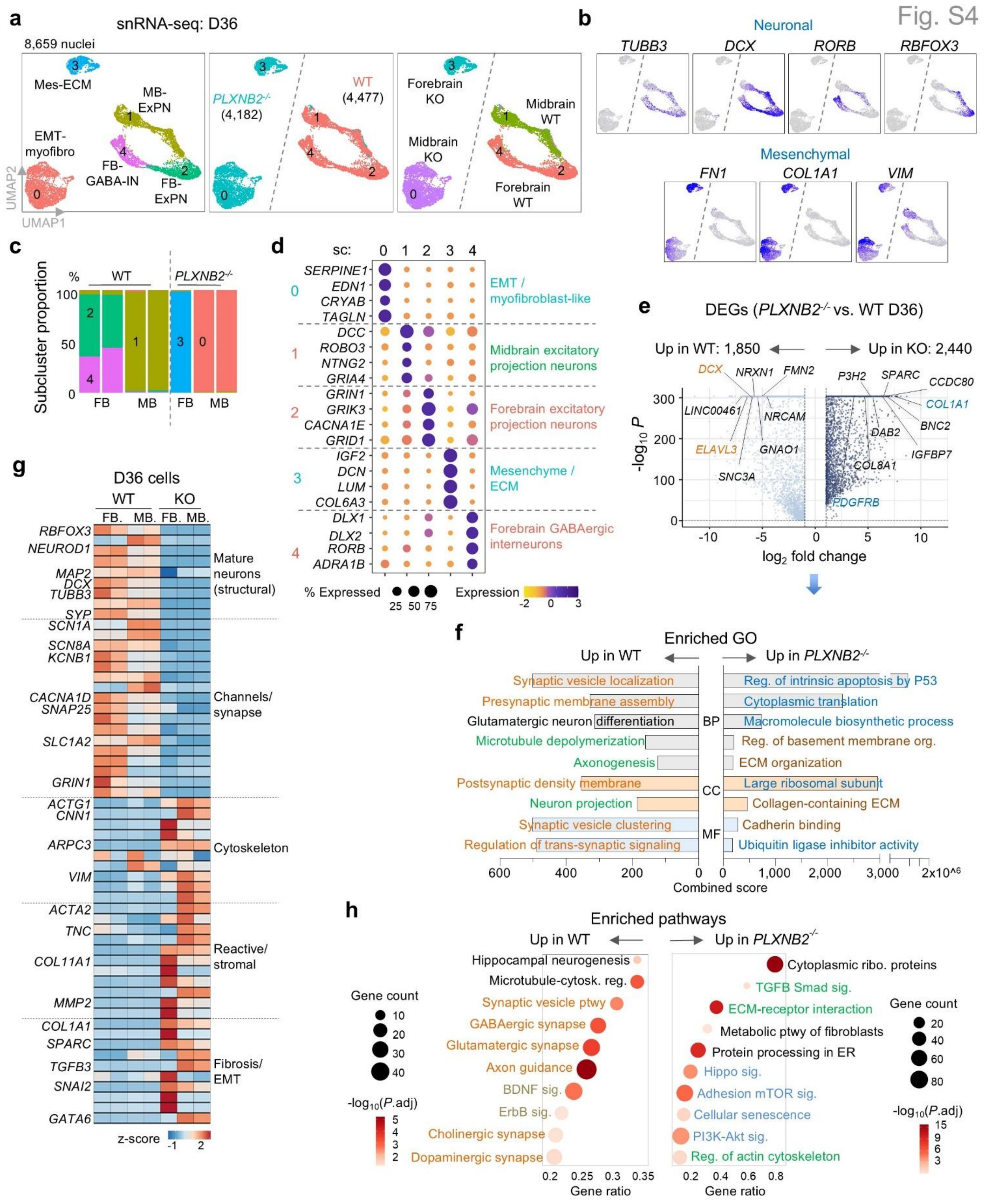

**Figure S4. Mismatched extrinsic cues lead to lineage instability.**

**a)** UMAP embedding of snRNA-seq data of D36 cells reveals major cell populations separated by genotype and by forebrain (FB) versus midbrain (MB) differentiation protocols.

**b)** Feature plots showing expression of representative marker genes for subclusters.

**c)** Stacked bar graphs showing relative proportions of subclusters in D36 WT and *PLXNB2*$^{-/-}$ cells across both FB and MB differentiation protocols. Graphs derived from seven combined snRNA-seq samples (two WT-FB, two WT-MB, and one KO-FB and two KO-MB).

**d)** Dot plot showing expression of marker genes in subclusters of D36 cells, with major themes annotated on the right and color-coded according to genotypes and FB vs. MB protocols.

**e)** Volcano plot of differentially expressed genes (DEGs) in *PLXNB2*$^{-/-}$ versus WT cells (pooled across FB and MB protocols), with top genes highlighted.

**f)** Gene ontology (GO) enrichment analysis of DEGs, grouped by biological process (BP), cellular component (CC), and molecular function (MF), color-coded by theme.

**g)** Heatmap of selected DEGs grouped by function in D36 WT and *PLXNB2*$^{-/-}$ cells cultured with FB or MB protocols.

**h)** Bubble plot showing enriched pathways upregulated in D36 *PLXNB2*$^{-/-}$ versus WT cells.

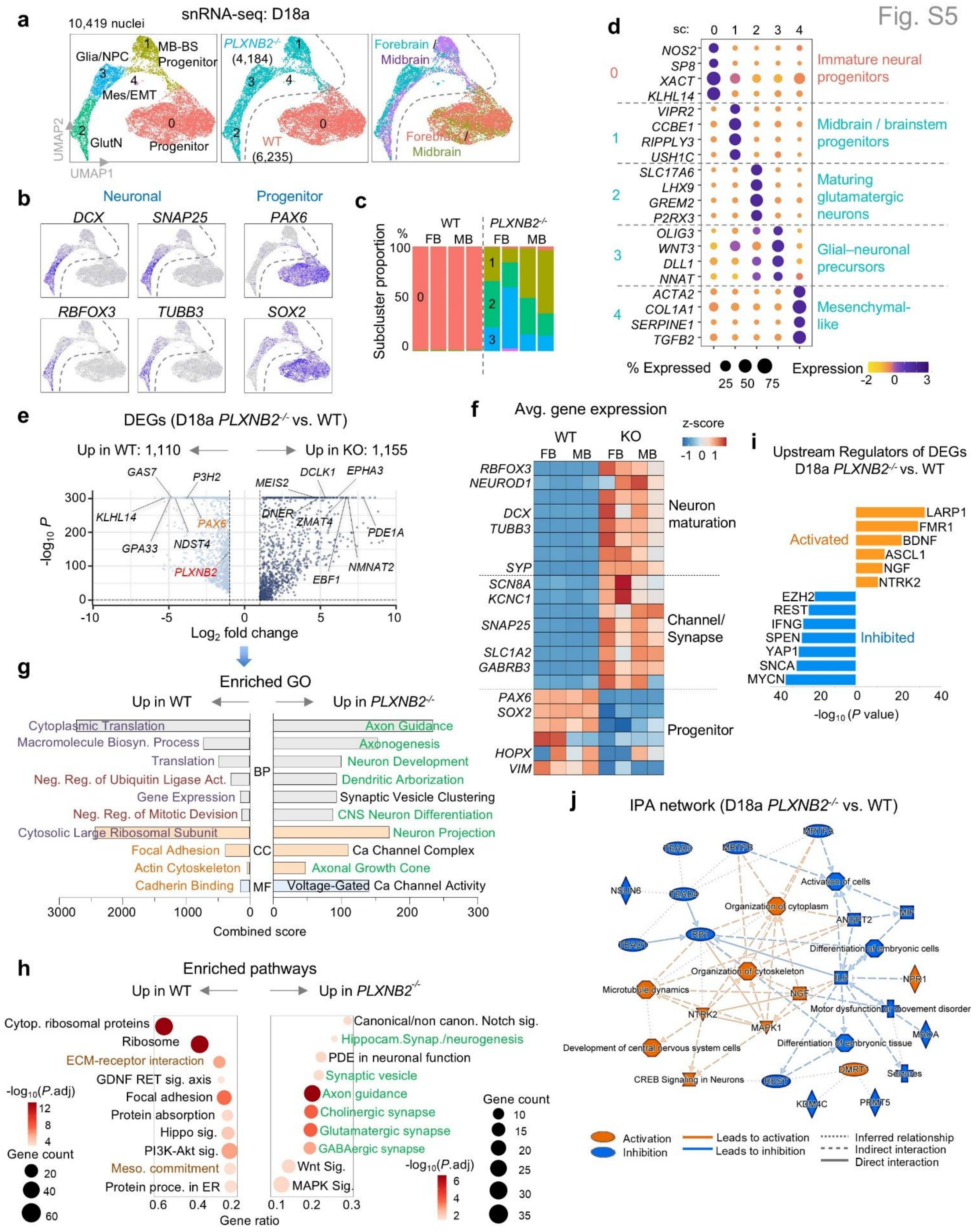

**Figure S5. Accelerated maturation cues stabilize neuronal lineage of PLXNB2$^{-/-}$ iNs.**

**a)** UMAP embedding of D18a cells under the accelerated protocol (switch to maturation media after the first passage at D9). WT cells (forebrain and midbrain protocols) clustered together as sc-0, whereas *PLXNB2*$^{-/-}$ cells segregated into four distinct subclusters.

**b)** Feature plots showing expression of representative marker genes of each subcluster.

**c)** Bar graphs illustrating distinct cell type proportions for WT and *PLXNB2*$^{-/-}$ cells under forebrain (FB) or midbrain (MB) iN protocols.

**d)** Dot plot of marker gene expression across subclusters of D18a cells.

**e)** Volcano plot of DEGs (*PLXNB2*$^{-/-}$ vs. WT) at D18a (pooled across FB and MB protocols), with top DEGs labeled.

**f)** Heatmap of representative DEGs grouped by functional categories, comparing WT and *PLXNB2*$^{-/-}$ cells under FB and MB protocols.

**g)** GO enrichment analysis of DEGs, grouped into biological process (BP), cellular component (CC), and molecular function (MF) categories, color-coded by theme.

**h)** Bubble plot showing pathways significantly enriched in *PLXNB2*$^{-/-}$ versus WT cells.

**i)** Upstream regulator analysis predicts activation of neuronal regulators (BDNF, ASCL1, NGF) and inhibition of progenitor regulators (REST, EZH2, YAP1) in *PLXNB2*$^{-/-}$ D18a cells.

**j)** IPA network analysis of *PLXNB2*$^{-/-}$ versus WT cells at D18a reveals activation of cytoskeletal remodeling, CREB signaling, and neuronal development, alongside suppression of progenitor maintenance and proliferation pathways.

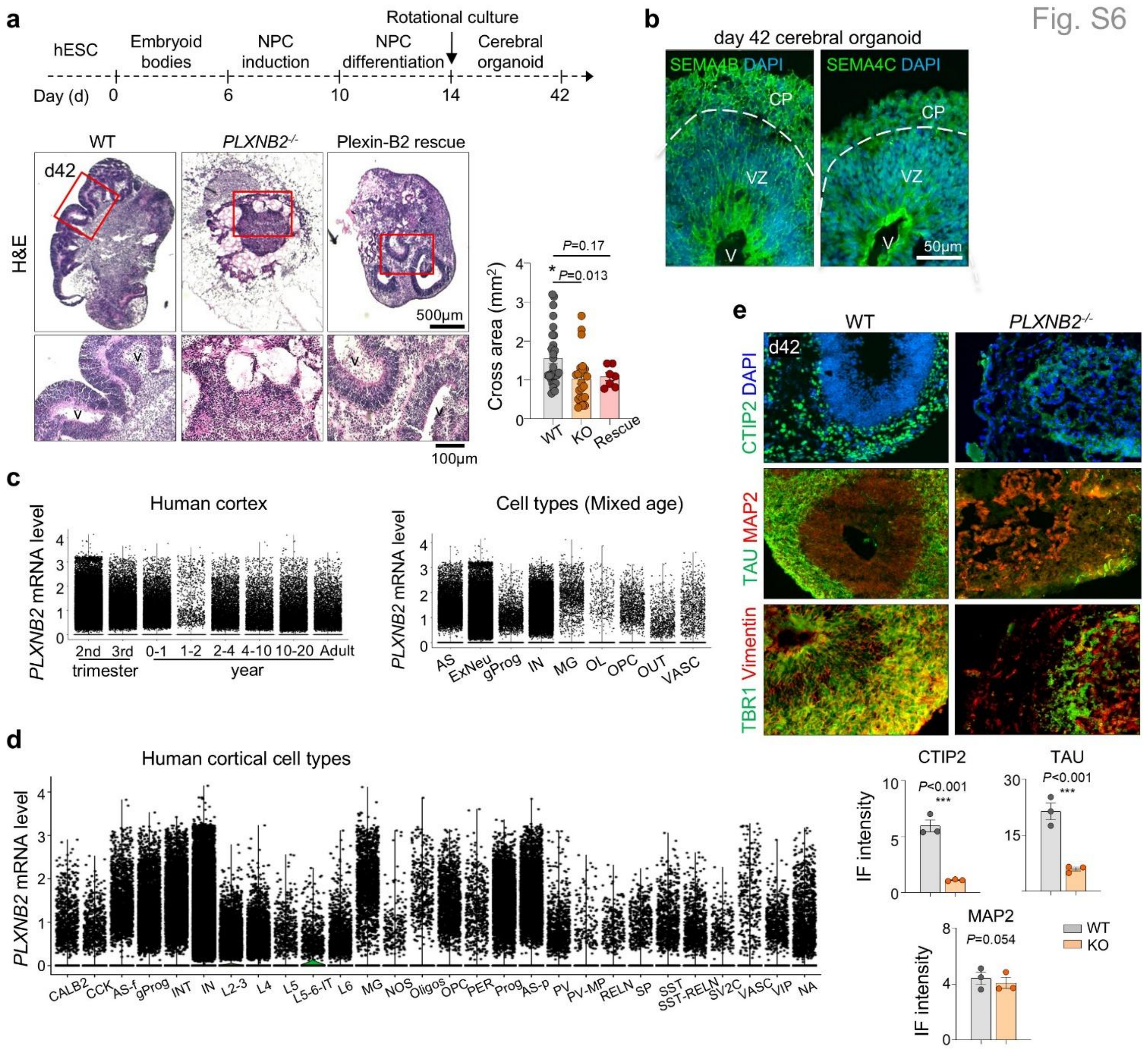

**Figure S6. Plexin-B2 in required for cortical structure during neurogenesis.**

**a)** Top, schematic of the cerebral organoid protocol. Bottom, H&E staining (enlarged images at bottom panel) showing that WT organoid at day 42 contained organized cortical architecture with ventricle-like (V) structures surrounded by ventricular zone (VZ) and cortical plate (CP). In contrast, *PLXNB2*$^{-/-}$ organoids displayed aberrant cystic morphology lacking VZ organization. Re-expression of Plexin-B2 rescued the KO phenotype. Quantification of organoid cross-sectional area, n = 6-14 organoids per group, one-way ANOVA with Dunnett's multiple comparisons test. Bar graphs show mean ± SEM.

**b)** Expression of SEMA4B and SEMA4C, cognate Plexin-B2 ligands, in d42 WT cerebral organoids.

DAPI used for nuclear counterstaining.

**c)** Left, temporal *PLXNB2* expression levels across developmental stages in human cortex. Right, *PLXNB2* expression across major cortical cell classes, including astrocytes (AS), excitatory neurons (ExNeu), glial progenitors (gProg), interneurons (IN), microglia (MG), oligodendrocytes (OL), oligodendrocyte progenitor cells (OPC), outer radial glia (oRG), and vascular cells (VASC). Based on data from Velmeshev et al., 2023.

**d)** Expression of *PLXNB2* in defined neuronal and non-neuronal subtypes of the developing human cortex, including excitatory neurons (L2–L6, L5–6 IT), inhibitory neuron classes (PV, SST, VIP), radial glia, astrocyte populations (AS-f, AS-p), microglia (MG), oligodendroglia (OPC, OL), and non-neuronal populations (VASC, NOS, NA). Based on data from Velmeshev et al., 2023.

**e)** *PLXNB2*$^{-/-}$ d42 organoids contained fewer CTIP2$^{+}$ deep-layer cortical neurons, which were arranged in disorganized manner. Additional markers MAP2 (dendrite marker), TAU (axon marker), vimentin (radial glial scaffold marker), and TBR1 (layer VI cortical neuron marker) reveal disrupted cortical lamination in *PLXNB2*-deficient organoids. Quantification based on n = 3 independent organoids per condition, with ~30 fields of view analyzed per organoid. Bar graphs represent mean ± SEM; two-tailed nested t-test.

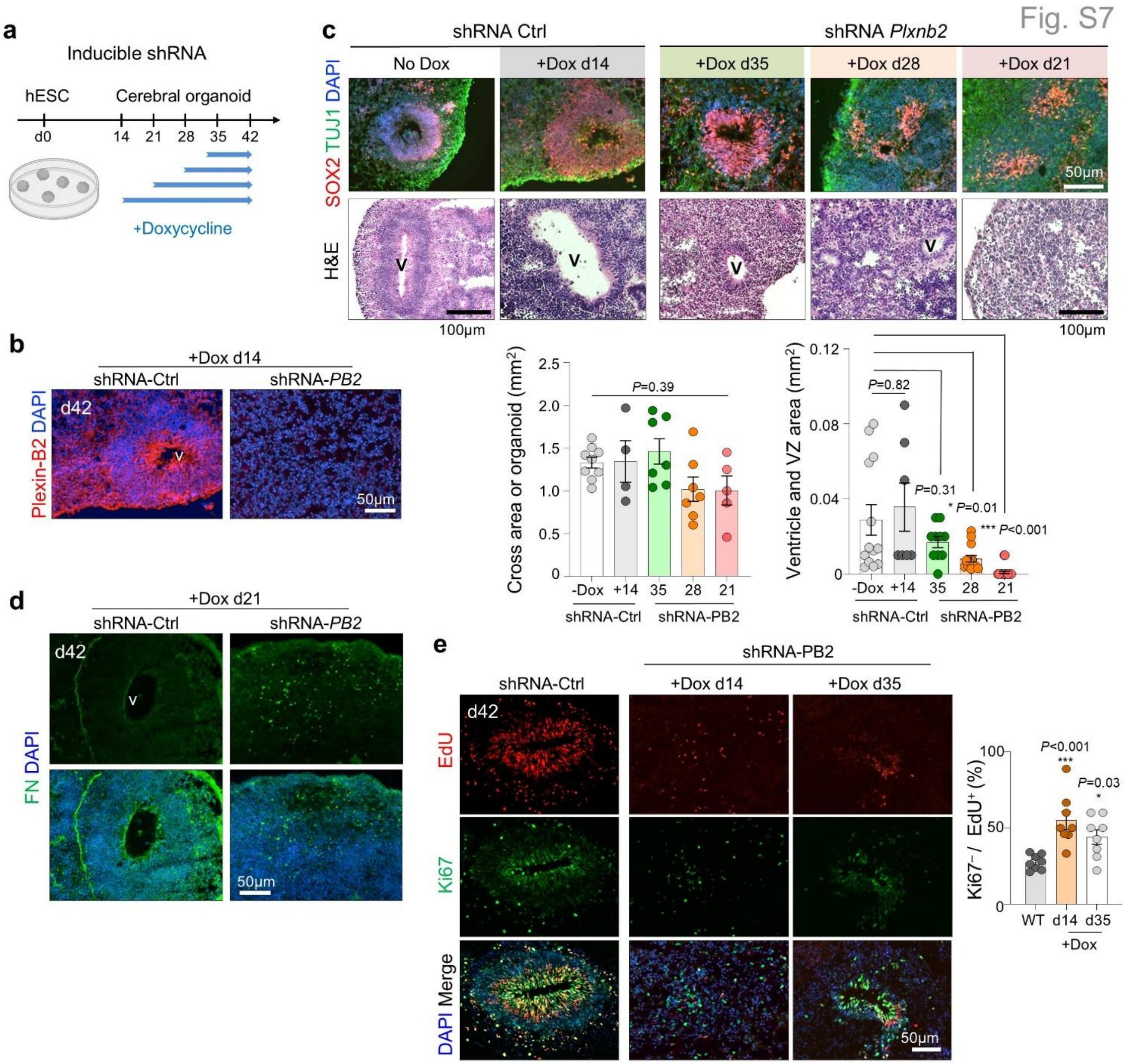

**Figure S7. *PLXNB2* deficiency disrupts cerebral organoid development.**

**a)** Experimental scheme of doxycycline (Dox)-inducible *PLXNB2* shRNA knockdown (KD) initiated at different timepoints.

**b)** IF confirming loss of Plexin-B2 protein expression in d42 cerebral organoids after KD induction at day 14.

**c)** Temporal KD analysis showing that earlier KD (day 21) caused more severe structural disorganization compared to KD at later stages. IF for SOX2 (progenitors) and TUJ1 (neurons), together with H&E staining, demonstrate progressive loss of ventricular morphology and progenitor zone integrity. n = 4-

12 organoids per group, with two randomly selected regions per organoid. One-way ANOVA with Dunnett's post hoc test. Bar graphs show mean ± SEM.

**d)** IF for fibronectin (FN) at d42 showing disruption of the FN network and abnormal ventricle-like structures in *PLXNB2* KD organoids.

**e)** Pulse-chase proliferation assay in *PLXNB2* KD cerebral organoids with EdU and Ki67 staining. Both early (d14) and later (d35) KD led to disrupted ventricular morphology and premature cell cycle exit, with earlier KD causing a more severe phenotype. n = 8 organoids per group, with two randomly selected regions per organoid. One-way ANOVA with Dunnett's multiple comparisons test. Bar graphs show mean ± SEM.

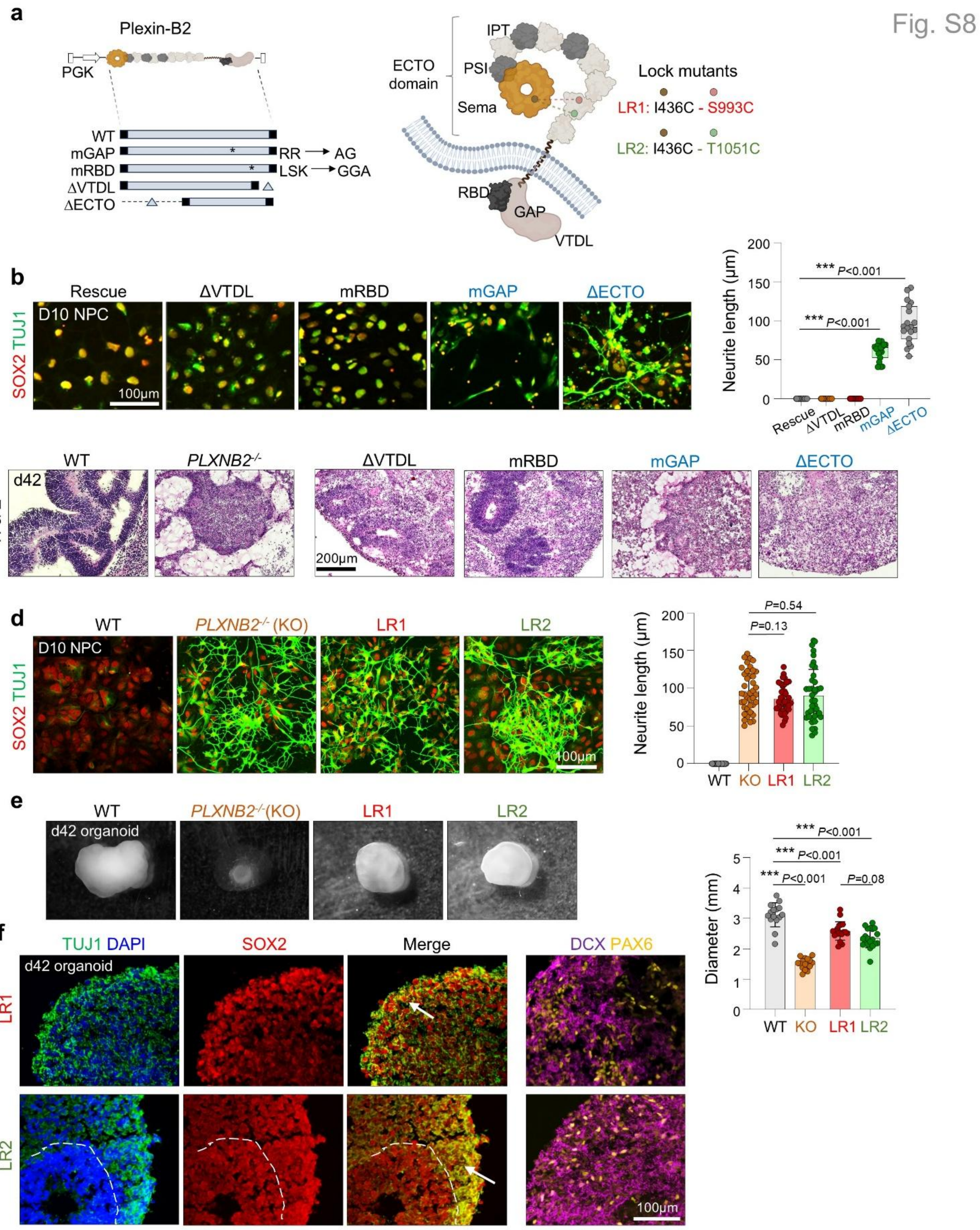

**Figure S8. Plexin-B2 mutational analysis reveals critical domains for gating neuronal differentiation.**

**a)** Left, mutations in specific functional domains of Plexin-B2. mGAP: mutated GTPase activating protein domain; mRBD: mutated Rho binding domain; ΔVTDL: deleted VTDL PDZ-binding motif at C-terminus; ΔECTO: deleted extracellular domain. Right: cartoon model indicating arrangement of Plexin-B2 domains. LR1 and LR2 indicated the "locked ring" mutations designed to prevent bending of the ECTO domain ring by mechanical stimuli.

**b)** Rescue experiments show that at D10 of iN protocol, mGAP and ΔECTO mutations failed to rescue the premature differentiation phenotype of Plexin-B2 KO. n=18 fields for each condition. One-way ANOVA with Dunnett's post hoc test. Box plots show median and quartiles.

**c)** Cerebral organoids carrying different domain-specific mutations of Plexin-B2 at day 42.

**d)** IF images of D10 cells in rescue experiments show that the locked ring mutations LR1 and LR2 phenocopy Plexin-B2 KO. n = 40 neurons for each condition. One-way ANOVA with Dunnett's post hoc test. Bar graphs show mean ± SEM.

**e)** Cerebral organoids derived from *PLXNB2*$^{-/-}$ or LR1 or LR2 mutant hESCs showed reduced size. Each data point represents one organoid (n = ~20 organoids per group). One-way ANOVA with Dunnett's multiple comparisons test. Bar graphs show mean ± SEM.

**f)** Cerebral organoids from LR1 or LR2 mutant hESCs at d42 show SOX2$^{+}$ progenitor cells overlapping with TUJ1$^{+}$ neurons (arrows), and disorganization of PAX6$^{+}$ and DCX$^{+}$ populations.

**Table S1. Overview of single nucleus RNA sequencing samples.**

| Protocol | Day of culture | Sample ID | Cell numbers (post-QC) |
|---|---|---|---|
| **Induced neurons** | **D9** | WT-1 | 889 |
| | | WT-2 | 2,595 |
| | | WT-3 | 1,490 |
| | | | *total 4,974* |
| | | KO-1 | 564 |
| | | KO-2 | 1,007 |
| | | KO-3 | 1,255 |
| | | | *total 2,826* |

| | | | |
|---|---|---|---|
| **Induced neurons** | **D36** | WT1 - Forebrain | 1,339 |
| | | WT2 - Forebrain | 1,173 |
| | | WT1 - Midbrain | 960 |
| | | WT2 - Midbrain | 1,005 |
| | | | *total 4,477* |
| | | KO1 - Forebrain | 1,027 |
| | | KO1 - Midbrain | 1,489 |
| | | KO2 - Midbrain | 1,666 |
| | | | *total 4,182* |

| | | | |
|---|---|---|---|
| **Induced neurons -** | **D18a** | WT1 - Forebrain | 1,890 |
| **Accelerated protocol** | | WT2 - Forebrain | 1,525 |
| | | WT1 - Midbrain | 1,451 |
| | | WT2 - Midbrain | 1,369 |
| | | | *total 6,235* |
| | | KO1 - Forebrain | 632 |
| | | KO2 - Forebrain | 1,273 |
| | | KO1 - Midbrain | 1,341 |
| | | KO2 - Midbrain | 938 |
| | | | *total 4,184* |

| | | | |
|---|---|---|---|
| **Cerebral organoid** | **d42** | WT-1 | 2,600 |
| | | WT-2 | 3,181 |
| | | WT-3 | 2,141 |
| | | | *total 7,922* |
| | | KO-1 | 4,955 |
| | | KO-2 | 211 |
| | | KO-3 | 1,853 |
| | | | *total 7,019* |

**Supplementary File 1**
Halperin et al.

**Molecular dynamics simulations**

Our recent investigations have advanced the understanding of how fundamental biophysical mechanisms govern cellular behavior, particularly focusing on the guidance receptor Plexin-B2. In references (Junqueira Alves et al. 2021; Junqueira Alves et al. 2025) we have studied Plexin-B2's critical role in regulating membrane tension and mechano-electrical coupling crucial for glioblastoma cell invasion through confined space and explored Plexin-B2's orchestration of actomyosin contractility, cytoskeletal tension, and adhesion during the collective dynamics of human embryonic stem cells and neuroprogenitor cells. Both studies utilized molecular dynamics simulations based on a coarse-grained bead model, representing the cell plasma membrane, nuclear membrane, and actin filaments.

Building upon this established framework, our current research aims to unravel the intricate cytoskeletal rearrangements underlying cell differentiation, specifically the morphological transformation of a stem cell into a neuronal cell. A key development in our model is the explicit inclusion of a microtubule component. This microtubule's growth rate is posited as a primary driver for the cell's morphological change.

The number of beads in a typical cell model is of order of N=700, which means that all the mass of the cell is homogenous distributed in order that $m_i = M_{cell}/N$, for a typical cell initial area $A_{cell} = \pi\, r^2{}_{cell}$ . The nuclear membrane is connected to the cell membrane through actin filaments-head interactions. Connections between the beads are modelled in a very special way, where springs of elastic potential with and long range interactions potentials are defined to model the main cell structures and morphologies.

To compute the forces of interactions between a given particle *i* and its neighbors, we chose some common potentials in the literature on the study of polymers. The potentials used in the model are Lennard-Jones Potentials ($U_{LJ}$), Weeks-Chandler-Anderson Potentials ($U_{WCA}$), Finite Extensible Nonlinear Elastic ($U_{FENE}$), and Bend Potential ($U_{Bend}$). The last allows the twisting of membranes, actin filaments and microtubules. Thus, the total potential to which each particle *i* is subjected is given by:

$$U_{tot}\left(\{\vec{r}\}, t\right) = U_{LJ}\left(\{\vec{r}\}, t\right) + U_{WCA}\left(\{\vec{r}\}, t\right) + U_{FENE}\left(\{\vec{r}\}, t\right) + U_{Bend}(\{\theta\}) \quad (1)$$

where $\{\theta\}$ is the set of all angles formed between three particles and $\{\vec{r}\}$ represent the set of all positions of partícles in the simulation. With the expression of the potential, it is possible to obtain the force that is subject to particle *i* is given by

$$\vec{F_i} = -\vec{\nabla}_i U_{tot}\left(\{\vec{r}\}, t\right) \quad (2)$$

Each bead was associated with potentials, which produced an interaction force with neighboring beads. Thus, these potentials are responsible for generating the dynamics of the system. As we are differentiating the cell membrane (particle 0), actin (particle 1), nucleus membrane (particle 3), heads of actin filaments (particle 7), microtubules (particle 12) and microtubules heads (particle 20), the elastic constants associated with each particle may be different. Thus, $\kappa_{0,0}$ is the elastic constant that connects the beads within the membrane (particles 0 with 0), $\kappa_{20,0}$ is the constant that connects particle 20 with particle 0, and so forth. For the Lennard-Jones potentials, we define the parameter $U_3$ as the strength of the $U_{LJ}$ interaction between beads in the actin filament head and the membrane.

In our coarse-grained bead model, the growth of this microtubule is simulated by assuming a constant growth rate for the distance between its constituent beads over time. To effectively capture this dynamic extension and its influence on cellular morphology, we are implementing a simultaneous, time-dependent update of the equilibrium parameters within our established potential functions. Specifically, the Lennard-Jones constant, denoted as $\sigma$, and the FENE equilibrium distance, $r_0$, are now dynamically modified. These parameters increase at a constant velocity over time until they reach a predefined maximum size, $r_{max}$. This methodological approach allows us to precisely simulate the continuous, directional force exerted by the growing microtubule, enabling the cell to undergo the intricate shape changes characteristic of neuronal differentiation, such as axon and dendrite extension. This advancement enhances the predictive nature of our models regarding mechanotransduction in developmental processes.

We observed that the interactions between the newly incorporated nanotube structure and the existing cellular components led to dynamic behaviors that closely matched those observed in experimental studies.

However, visual comparison of simulation snapshots with experimental images is often qualitative and insufficient for robust analysis. To provide a quantitative measure of this morphological development, we created a method to classify and quantify the extent to which the simulated protuberance resembles a neuron's growth pattern, which we call the Area Metric. First of all, we need to transform all positions into the nucleus center's frame of reference, which differs from the center of mass, by $\vec{r} = \vec{r}' - \vec{r}_{CN}$, where $\vec{r}'$ is the position of the membrane structure, $\vec{r}_{CN}$ is the nucleus center and $\vec{r}$ the relative positions. To analyze the spatial characteristics of the membrane structures more effectively, the data was transformed into polar coordinates. Unlike cartesian coordinates, which describe positions in terms of linear distances along fixed axes, polar coordinates allow us to express points in terms of their radial distance from a reference point and their angular displacement.

By using polar coordinates, we can more easily quantify radial variations, identify directional patterns, and distinguish growth trends. This approach provides a way to describe shape deformations and facilitates the comparison of cells with different orientations or sizes.

Now, we can establish a criterion to define what constitutes a protuberance using the outlier method (SciPy Community). To achieve this, we calculate the standard deviation and mean radius for each frame of our simulation and defined a limit $\lambda = r_M + \frac{3}{2}\sigma_M$, where $r_M$ is the mean radius, $\sigma_M$ the variance.

With the outliers identified, we can now calculate the area of the graph above the defined limit using integration methods:

$$Area = \frac{1}{2}\int_{\theta_1}^{\theta_2} (r(\theta))^2 \, d\theta \qquad (3)$$

We can now calculate the area of the graph above the defined limit using numerical integration, specifically the trapezium (trapezoidal) method (SciPy Community). This approach is well-suited for discrete data points, as it approximates the area under a curve by dividing it into a series of adjacent trapezoids.

This method is efficient because it accounts for the entire membrane while distinguishing true protuberances from irregular or asymmetric growth. In cases where the cell exhibits asymmetric expansion or minimal overall growth, the computed area remains low, as deviations from the mean radius are distributed rather than concentrated. Otherwise, when a single prominent protuberance develops, the outlier method effectively captures its extent, resulting in a high area score. This selective sensitivity makes the method robust in identifying significant structural changes while minimizing false positives from minor fluctuations in membrane shape.

SciPy Community, "scipy.integrate.trapezoid," SciPy v1.15.2 Manual. [Online]. Available: https://docs.scipy.org/doc/scipy-1.15.2/reference/generated/scipy.integrate.trapezoid.html.
